# Tuning Insulin Receptor Signaling Using *De Novo* Designed Agonists

**DOI:** 10.1101/2024.10.07.617068

**Authors:** Xinru Wang, Sarah Cardoso, Kai Cai, Preetham Venkatesh, Albert Hung, Michelle Ng, Catherine Hall, Brian Coventry, David Lee, Rishabh Chowhan, Stacey Gerben, Jie Li, Weidong An, Mara Hon, Domenico Accili, Xiaochen Bai, Eunhee Choi, David Baker

## Abstract

Binding of insulin to the insulin receptor (IR) induces conformational changes in the extracellular portion of the receptor that lead to activation of the intracellular kinase domain and the AKT and MAPK pathways, and downstream modulation of glucose metabolism and cell proliferation. We reasoned that designed agonists that induce different conformational changes in the receptor might induce different downstream responses, which could be useful both therapeutically and to shed light on how extracellular conformation is coupled to intracellular signaling. We used *de novo* protein design to first generate binders to individual IR extracellular domains, and then to fuse these together in different orientations and with different conformational flexibility. We describe a series of synthetic agonists that signal through the IR that differ from insulin and from each other in the induction of receptor autophosphorylation, MAPK activation, intracellular trafficking, and cell proliferation. We identify designs that are more potent than insulin causing much longer lasting reductions in glucose levels, and that retain signaling activity on disease-causing receptor mutants that do not respond to insulin. These results inform our understanding of how changes in receptor conformation and dynamics are transmitted to downstream signaling, and our synthetic agonists have considerable therapeutic potential for diabetes and severe insulin resistance syndromes.

**Highlights:**

- Computational design yielded super agonists, partial agonists, and antagonists of IR.
- *De novo* agonists induce a distinct IR active conformation.
- Designed agonists tune IR signaling by modulating conformational dynamics of activated IR.
- Designed agonists are more potent than insulin, reducing glucose levels longer and activating disease-causing IR mutants.

## Introduction

Insulin receptor (IR), a receptor tyrosine kinase, plays an important role in metabolism, development, growth, and proliferation ^1–3^. Insulin activated IR undergoes trans-autophosphorylation and phosphorylates a number of intracellular substrates, triggering two major signaling pathways – protein kinase B (AKT) pathway and the MAP Kinase (MAPK) pathway ^4–7^. The phosphorylation cascades control trafficking events, transcription factors, glucose and lipid metabolism, and growth in different tissues and pathophysiological conditions. Dysregulation of IR signaling causes diseases including diabetes, cancer, and aging. IR is a dimer composed of two protomers linked by disulfide bonds. The structures of IR in inactive and active states have been determined (Fig. 1A) ^8–15^. Insulin binding at two distinct sites in the extracellular domains of IR promotes a conformational change of IR which reduces the distance between the intracellular kinase domains which leads to trans-autophosphorylation. The primary insulin binding site (site-1) is located in the L1/α-CT domains of IR, while the secondary insulin binding site (site-2) is located on the side of the F1 domain (Fig. 1A,B). It has been proposed that the conformational dynamics of IR in different ligand-bound active states are critical for their biological functions ^10,11^. However, because of the complexity of these large complexes, and the similarity in conformation observed for ligands that exert different effects on signaling, achieving a full understanding of how receptor conformational dynamics are linked to diverse downstream signals, trafficking, and biological outcomes is a current challenge.

**Figure 1.**
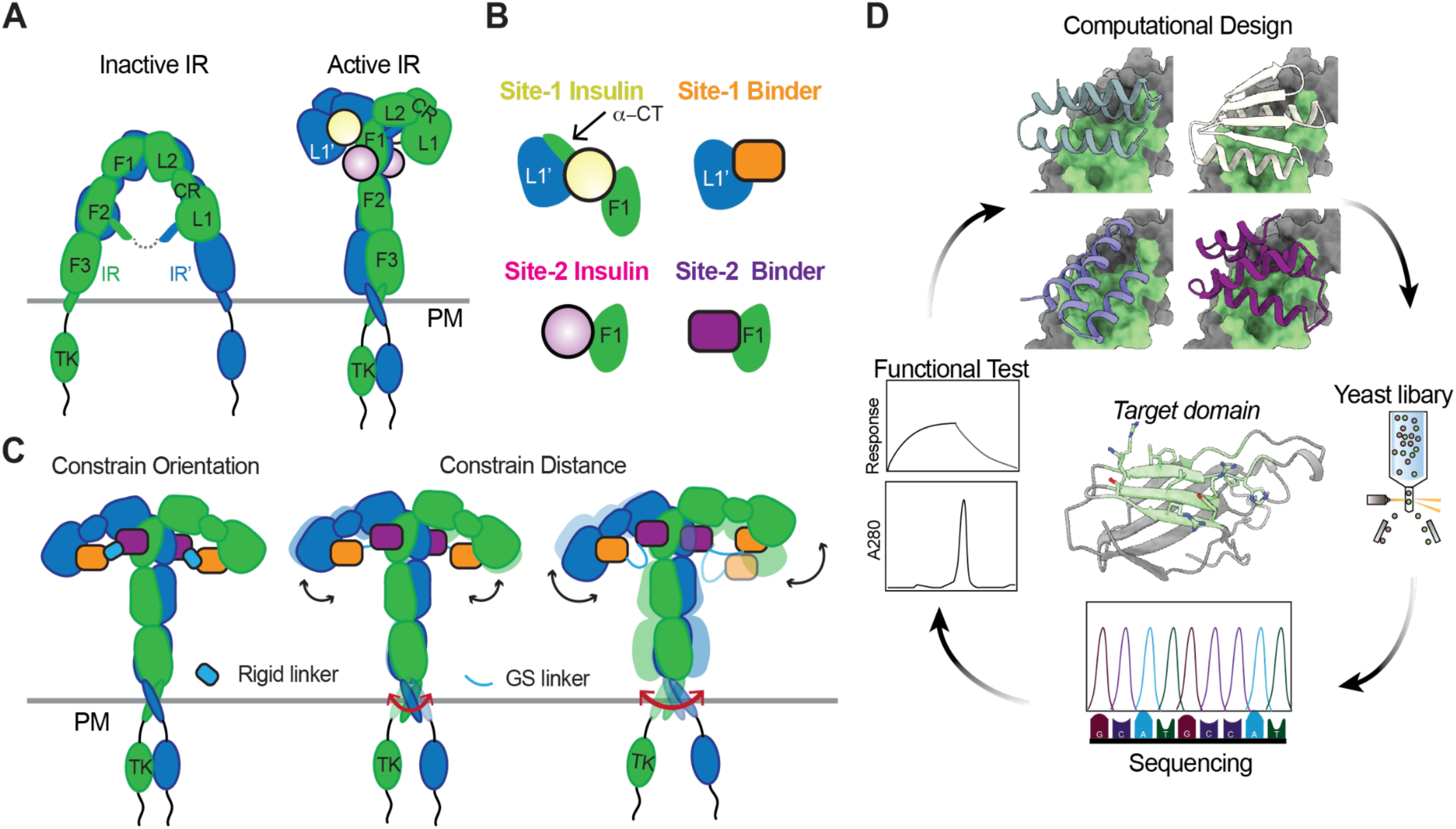
Insulin receptor agonist design strategy. (A) Schematic representation of the structure of the inactive, apo-IR and insulin-bound, active IR. Protomer 1 (green) and protomer 2 (blue). Leucine-rich repeats 1 and 2 (L1, L2), Cysteine-rich (CR), Fibronectin type III-1, 2, and 3 (F1, F2, and F3), C-terminal tail of α-subunit (α-CT), tyrosine kinase (TK) domains. Domains in protomers 1 and 2 are shown as L1-TK and L1’-TK’, respectively. Site-1 insulins (yellow) and Site-2 insulins (pink). (B) Close up view of the tripartite interaction between insulin and IR L1’, α-CT, and F1 domains (Site-1 insulin), and the interaction between insulin and IR F1 domain (Site-2 insulin) (left). Target domains and designer binders (right). (C) Design strategy to regulate IR conformations and dynamics by flexibly or rigidly fusing site-1 and site-2 binders. When S1B and S2B are fused rigidly, the relative positions of the L1 and F1 domains of IR are fixed. In contrast, when they are fused with a flexible linker, the ligand-bound receptor should retain flexibility. (D) Scheme of protein binder development approach. Computationally designed protein binders will be screened using yeast display and sorted by FACS. The enriched binders will then be identified by NGS and further characterized experimentally. Representative designed IR site-2 binders are shown in the top panel.

We reasoned that *de novo* protein design could provide a solution to this problem. There are only a small number of natural ligands for any given receptor, and the conformational dynamics they produce in the target receptor are not easily modulated; in contrast protein design can in principle generate a wide range of ligands that bind to multiple domains in receptor extracellular regions and tune their relative orientations and dynamics. To probe the relationship between extracellular conformation of the IR and intracellular signaling, we set out to design proteins that modulate the conformational dynamics of the extracellular domains of IR and to determine the effect of these on the extent of induced autophosphorylation and downstream signaling (Fig. 1C). Such designed IR agonists could also have therapeutic potential: although recombinant insulin and its analogs have been used to treat type 1 and type 2 diabetes for nearly a century, these treatments are not without limitations, including complications in manufacturing processes and storage, which could potentially be reduced for hyperstable, easy-to-manufacture designed proteins^16^.

## Results

We hypothesized that synthetic molecules that engage two insulin binding sites in the extracellular domains of IR and induce conformational changes to reduce the distance between intracellular kinase domains could activate IR signaling. To generate new IR agonists that bring together the different domains in the extracellular region in different orientations, we used a two-step approach: we first sought to design binders for the L1 domain and F1 domain of IR (Fig. 1B) and second to fuse the individual binders together to induce conformational rearrangements in the receptor (Fig. 1C). Such synthetic binders could break the autoinhibitory conformation of IR by displacing the α-CT motif from the L1 domain of IR, and stabilize the active conformation by simultaneously binding to two protomers. We reasoned that by varying the rigidity of the linker between the two binding domains, we could explore the effects of the conformational stability of the active state of IR on downstream signaling, intracellular trafficking, and its functions (Fig. 1C). Previously, a miniprotein binder was developed to bind the L1 domain of IR using a Rosetta based design pipeline ^17^ (we refer to this below as S1B). Therefore, we started by designing IR site-2 binders.

### *De novo* design of IR site-2 binders

We set out to design site-2 binders that bind to the outer side of the F1 domain in the inactive state of IR (Fig. 1B,D). We used the Rosetta RIF dock method^17^ followed by Rosetta Fast Design to design binders to the F1 domain structure from a high-resolution cryo-EM model (PDB:6PXV)^8,17^, and filtered the designs based on Rosetta metrics (‘ddg’, ‘sasa’, ‘contact_molecular_surface’,’contact patch’)^17^ and DeepAccNet (‘Plddt’)^18^. We constructed a library of 11,452 designs that passed the filters and screened them using yeast display (Fig. 1D). Seventeen designs bound the F1 domain on the yeast surface at 1 nM (Fig. S1A,B). After optimization through site-saturation mutagenesis (SSM) and combination mutagenesis, we selected several designs and determined their binding affinity for the F1 domain of IR to be sub-nanomolar to nanomolar using biolayer interferometry (BLI). These binders are either helical bundles or ferredoxin scaffolds (Fig. 2A, Fig. S1B) and are modeled to bind to insulin binding sites on the F1 domain. One binder, referred to as S2B, which binds to the F1 domain of IR with a *K_D_* of 1.9 nM, significantly higher than insulin (*K_D_* of 21 μM, Fig. S1D), was selected for further study (Fig. 2C). S2B is stable at 95 °C (Fig. 2D), whereas insulin irreversibly unfolds at high temperatures (Fig. S1C). We investigated the effect of the S2B design on IR signaling. There are two IR isoforms, a short isoform, IR-A and a long isoform, IR-B^19^. We treated DKO-IR-B cells (IR and IGF1R double knockout preadipocytes expressing human IR-B) with insulin wild-type (WT), insulin ValA3E (a site-1 binding defective mutant), and insulin LeuA13R (a site-2 binding defective mutant) in the presence and absence of S2B. Consistent with previous work, insulin ValA3E did not activate IR, while insulin LeuA13R partly activated IR and downstream signaling in DKO-IR-B cells (Fig. 2E,F). Cotreatment with insulin site-1 and site-2 binding defective mutants can activate IR^10^. Similarly, when the designed S2B was co-treated with the insulin site-2 binding defective mutant, we observed increased IR autophosphorylation (pY1150,1151 IR) and downstream signaling compared to insulin site-2 binding defective mutant or S2B alone (Fig. 2E,F). Similar results were obtained in cells expressing IR-A, a short isoform of IR (Fig. S1E,F). Thus, while inactive on its own, S2B can synergize with site-1 only binding insulin to activate both isoforms of IR.

**Figure 2.**
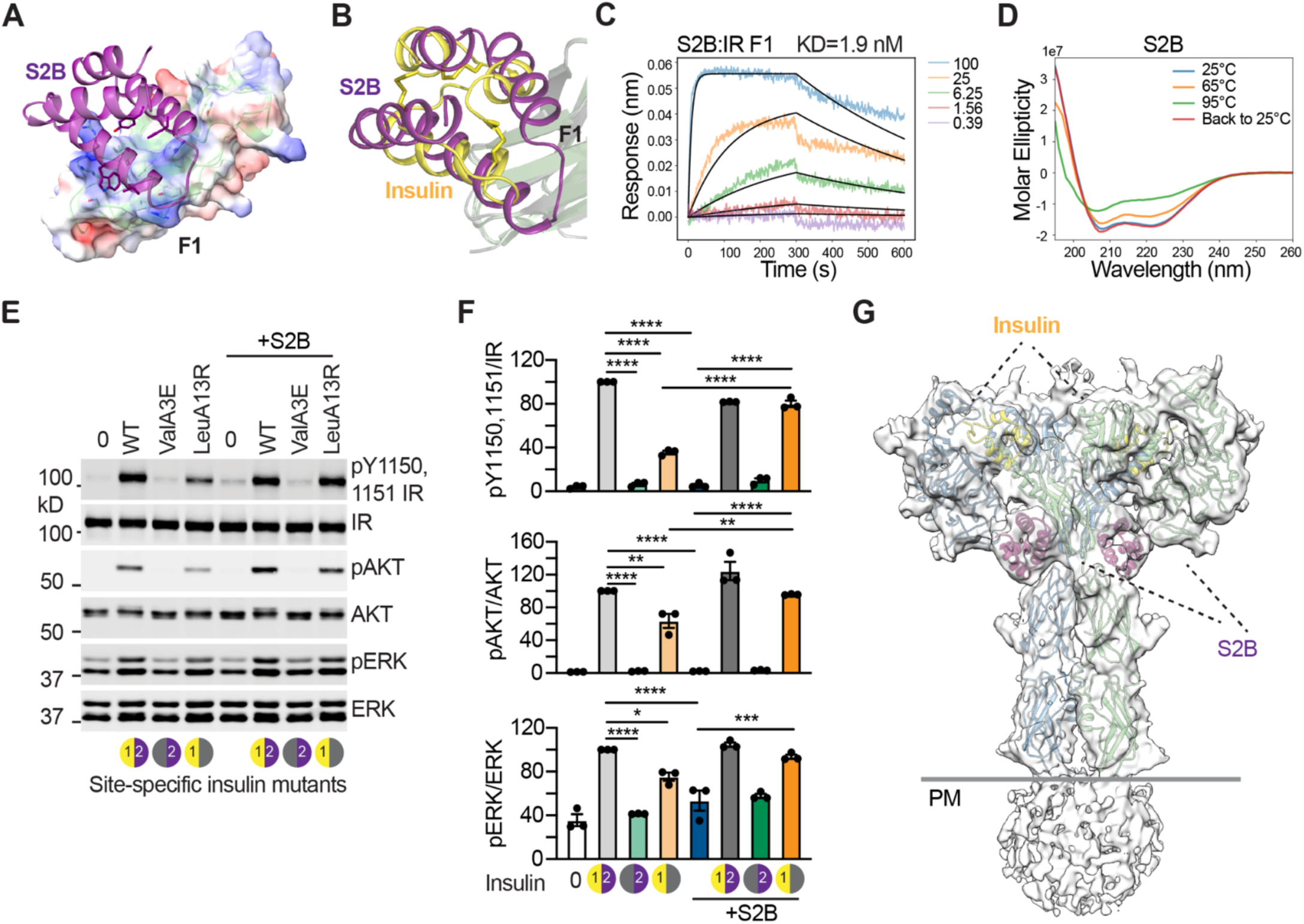
Characterization of IR site-2 binder. (A) Design model of S2B:IR F1 domain complex. The F1 domain is shown as transparent surface colored according to electrostatic potential and S2B is illustrated as purple cartons with key interaction residue side chains shown as sticks. (B) Superposition of S2B:IR F1 domain complex design model and insulin:F1 complex (PDB:6PXV). S2B binds to IR F1 domain at similar areas as insulin with different topology. (C) Biolayer interferometry characterization of binding of S2B to IR F1 domain. Global kinetic fit was reported. (D) CD spectra of S2B at various temperatures. S2B slightly unfolds at 95 °C but refolded when the temperature drops. (E) IR signaling in DKO-IR-B cells treated with 10 nM wild-type (WT) or site-specific insulin mutants with or without 100 nM site-2 binder (S2B) for 10 minutes. (F) Quantification of the western blot data shown in (E). Levels of autophosphorylation were normalized to total IR levels and shown as intensities relative to that in WT insulin alone. Mean ± sem. N=3 independent experiments. Significance calculated using two-tailed student’s t-test. (G) Cryo-EM model of Insulin/S2B/IR complex. Two protomers are shown in green and blue. Insulin and S2B are shown in yellow and purple cartoons. The cryo-EM density is shown as a transparent surface. PM, plasma membrane.

To determine the mechanism of receptor activation by S2B, we determined a 6 Å cryo-EM structure of IR bound with S2B together with insulin (Fig. 2G, S2). In the structure, two insulin and two designed S2B molecules bind to IR site-1 and site-2, respectively, and promote the symmetric T-shaped IR conformation. Thus, the functionally mimicry of site-2 binding insulin by S2B arises from similar effects on IR conformation.

### Design of Site-1 and Site-2 binder fusions

We next sought to link S2B with S1B (Fig. 3A,B) such that binding to the L1 and F1’ domains would break the autoinhibitory conformation, induce conformational rearrangement, and stabilize an active state of IR (Fig. 1C). We overlaid the design models of S2B/F1 and S1B/L1 on the active conformation of IR (PDB: 8DTL) by superimposing the F1 and L1 domains^20^. We then used RFdiffusion to build a rigid connector between the two binders^21^ (Fig. 3B,C). The first helix of L1B (Up to residue 3) and the last helix of S2B (Residue 54 onwards) were rebuilt by RFdiffusion to link the L1- and S2-binding interfaces. Surface residues near the diffused regions were masked and redesigned using ProteinMPNN along with the newly built regions. 32 sequences were generated for each backbone. Designs passing AF2^22^ and Rosetta metrics^17^ (pae interaction, plddt, ddg, contact surface) were selected for experimental characterization. To determine how conformational stability affects IR signaling, we also generated more flexible fusions by linking the two domains with GS linkers in both orientations; in the overlaid composite structure the distance between the S1B-C and the S2B-N terminus (S1-Fn-S2) is ∼18 Å and between the S2B-C terminus and S1B N-terminus (S2-Fn-S1) is ∼9 Å; we therefore used slightly longer linkers (Fn; n, linker lengths = 3 to 11 residues) in the S1-Fn-S2 than the S2-Fn-S1 constructs (Fn; n = 1 to 8 residues) (Fig. S6A).

**Figure 3.**
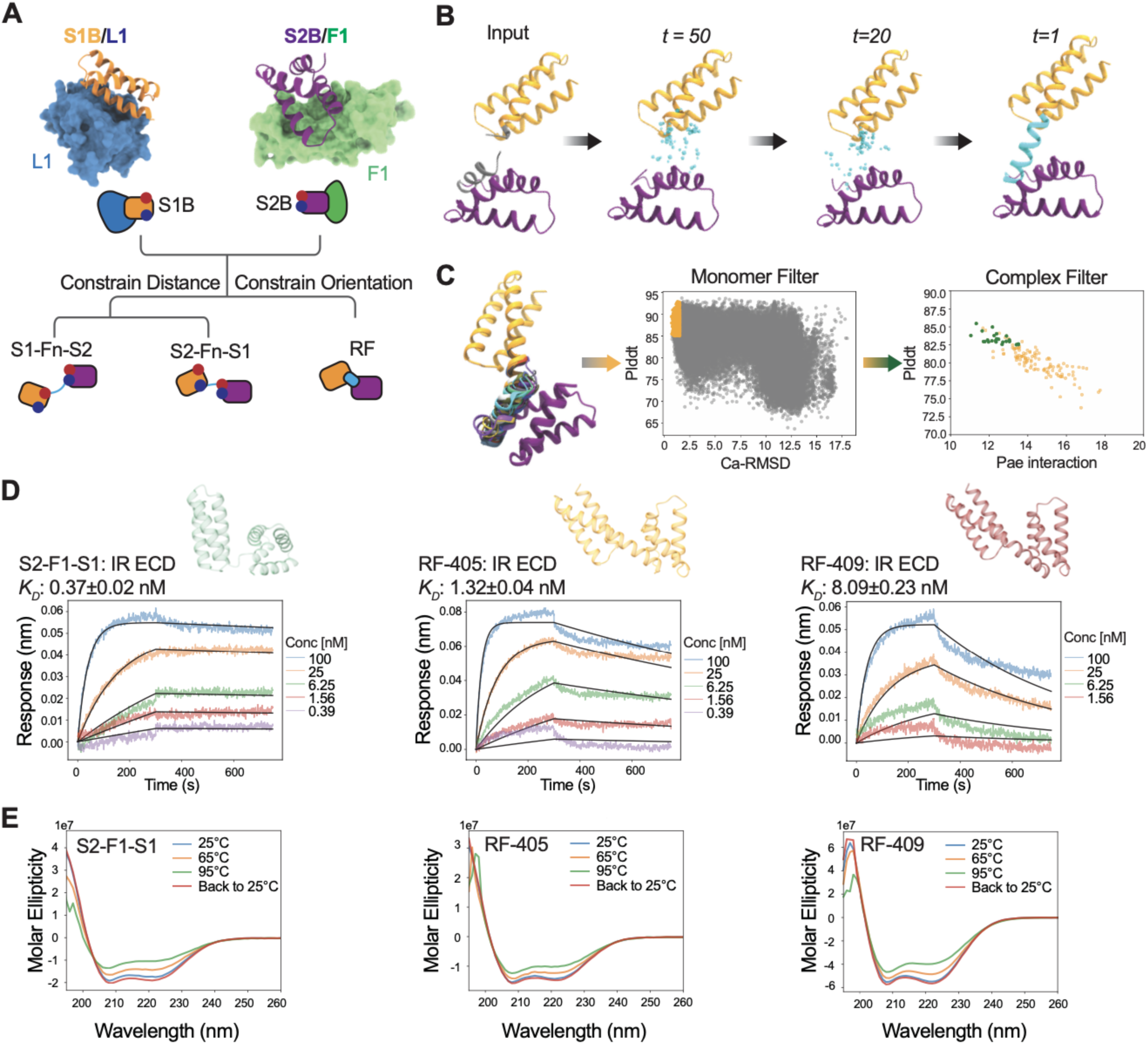
Develop *de novo* IR agonists. (A) Design of Site-1 and Site-2 binder fusions. To arrange L1’ domain and F1 domain into active conformation, S1B and S2B were fused with either flexible linkers or rigid interdomain connection. The N-terminus and C-terminus of S1B or S2B were highlighted with blue and red dots, respectively. (B) An example of design trajectory for building linker between the two binding domains. (C) RFdiffusion-generated backbones were designed using ProteinMPNN and predicted as monomers by AlphaFold2. These designs were aligned with the input structure, and those with a pLDDT score >85 and Cα-RMSD <1.5 were selected. The selected designs were then further predicted in complexes with the target PDB (L1+F1) and filtered again. (D) Design models and biolayer interferometry characterization of binding of S2-F1-S1, RF-405, and RF-409 to IR-ECD. Global kinetic fit was plotted. (E) CD spectra of S2-F1-S1, RF-405, and RF-409 at various temperatures.

The binding affinities of the two-domain binding constructs were estimated by BLI against the biotinylated IR extracellular domain (IR-ECD). S2-F1-S1 bound to IR-ECD with a *K_D_* of 0.37 nM, while RF-405 and RF-409 bound to IR-ECD with a *K_D_* of 1.3 nM and 8.1 nM, respectively (Fig. 3D). The fusion constructs are quite specific as they did not bind IGF1R, a homologous receptor tyrosine kinase (RTK) that can be activated by insulin^23,24,25,26^ (Fig. S1G). Circular dichroism (CD) temperature melting studies showed that both the flexibly-linked and rigidly-linked binders are hyper-thermostable (Fig. 3E).

### Cryo-EM complex structure determination

To determine whether the synthetic fusion constructs induce the intended conformational changes, we determined cryo-EM structures of the RF-405/IR and S2-F1-S1/IR complexes at resolutions 4 Å and 8 Å, respectively (Fig. 4A-D, S3-5). The cryo-EM structure of RF-405/IR complex exhibits an extended T-shaped architecture. A strong density between the two protomers at the top part of the IR was observed and unequivocally assigned to RF-405. The design models of the S1B and S2B of the RF-405 fit well into the cryo-EM density as rigid bodies without further refinement; consistent with the models, the S1B binds the L1 domain of IR (site-1), while the S2B contacts a side surface of the F1 domain of the adjacent IR protomer (site-2) (Fig. 4E-G, Fig. S3I). S1B and S2B of RF-405 are linked through a continuous α-helix as predicted in the design model (Fig. 4A,B). Both site-1 and site-2 interfaces involve both hydrophobic and electrostatic interactions, and Trp65 of RF-405 is sandwiched between the S1B and S2B components, enhancing the rigidity of the design (Fig. 4G). The α-CT motif is displaced from the L1 domain upon the binding of the S1B of RF-405. RF-405 crosslink the two IR protomers by simultaneously contacting the site-1 and site-2, thereby stabilizing the extended T-shaped active conformation (Fig. 4A,B). This conformation is similar to that induced by S597^20^, suggesting that the designed binders and S597 induce similar conformational changes for receptor activation.

**Figure 4.**
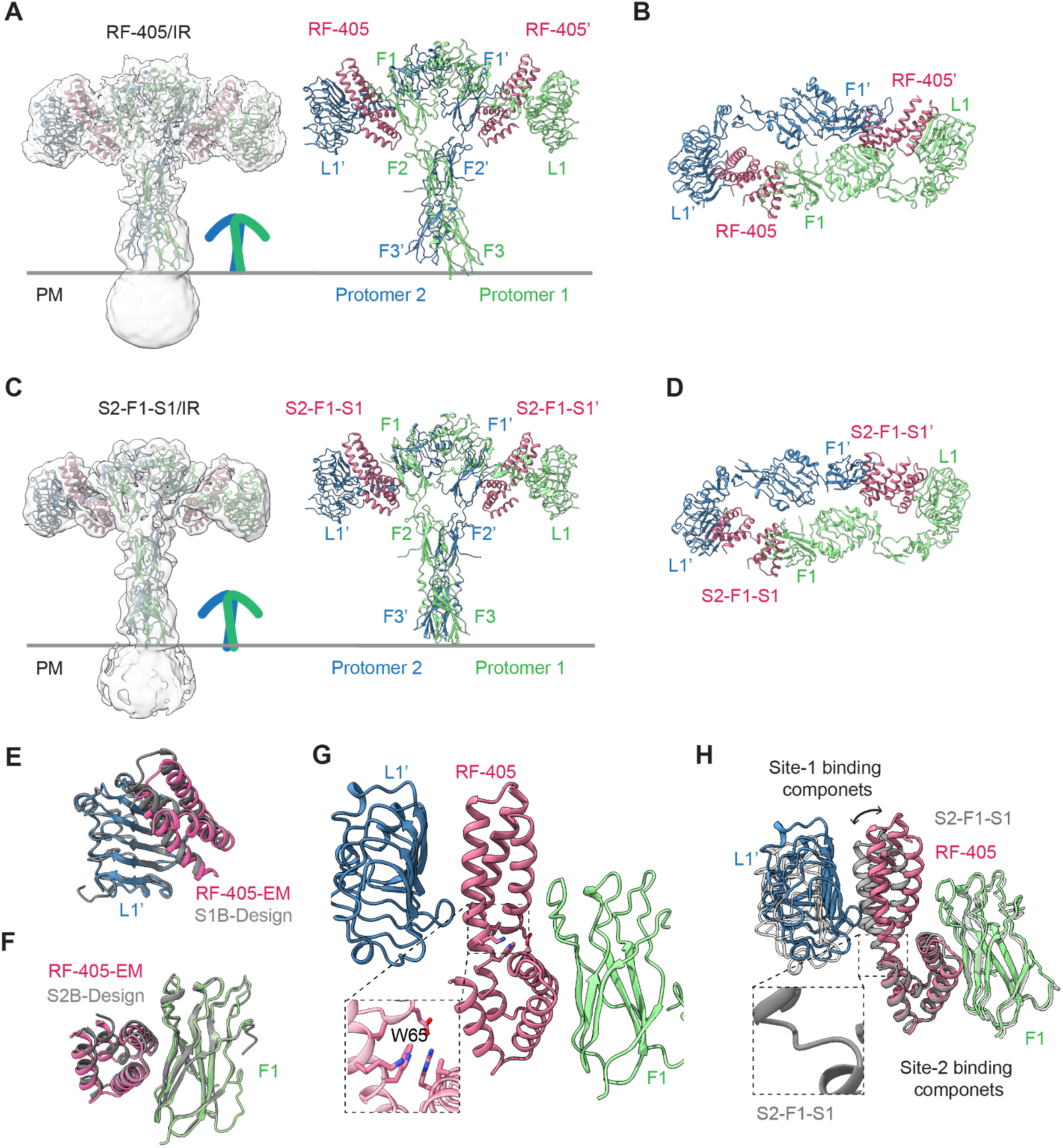
CryoEM structures of RF-405/IR and S2-F1-S1/IR complexes. (A) Cryo-EM model of RF-405/IR complex. Two protomers are shown in green and blue, RF-405 molecules are shown as pink cartoon. RF-405 molecules in each protomer are shown as RF-405 and RF-405’. The cryo-EM density is shown as a transparent surface. (B) Top view of RF-405/IR complex structure. (C) Overall structure of S2-F1-S1/IR complex. (D) Top view of S2-F1-S1/IR complex structure. (E) Close-up view of RF-405 (pink) binding at the L1 domain (blue) of one protomer. (F) Close-up view of RF-405 (pink) binding at the F1 domain (green) of one protomer. (G) Close-up view of the binding of RF-405 (pink) at the L1’ (blue) and F1 (green) domains of IR. Trp65 of RF-405 is sandwiched between the Site-1 and Site-2 binding components and highlighted. (H) Overlay of S2-F1-S1 (gray) and RF-405 (pink) aligned with the site-2 binding component. The linker of S2-F1-S1 is highlighted.

The cryo-EM structure of the S2-F1-S1 in complex with IR was resolved at a lower resolution than the RF-405/IR complex: the flexibility of S2-F1-S1 likely translates to increased flexibility of the complex relative to the rigid RF-405 (Fig. 4C,D, Fig. S3G). The structure of the S2-F1-S1/IR complex has a similar extended T-shape to that of RF-405/IR complex (Fig. 4C,D). S1B and S2B of S2-F1-S1 are linked by a short flexible loop (Fig. 4D,H); superimposition of the bound S2-F1-S1 and RF-405 revealed that, due to the rigid linkage, the distance between the S1B and S2B domains of RF-405 is shorter than those of S2-F1-S1 (Fig. 4H). The compact conformation of RF-405 allows its S1B component to simultaneously contact the L1 domain of one protomer and a loop in the top region of the F1 domain of the other protomer, further increasing the stability of the active conformation of IR.

We next attempted to determine the cryo-EM structures of the S1-F8-S2/IR complexes (Fig. S3J). Unlike S2B-S1B fusion constructs, 2D class averaging of S1-F8-S2/IR revealed a high degree of conformational heterogeneity. Even though individual domains could be identified in a subset of 2D class averages, no high-resolution features were apparent, such as clearly identifiable secondary structural elements. These data suggest that the S1-F8-S2/IR complex samples a much greater range of conformations than the S2-F1-S1 and RF-405 complexes.

### Characterization of synthetic agonists

We next tested the effects of the fusion constructs on IR activation and downstream signaling. S1-Fn-S2 increased levels of IR autophosphorylation (pY1150,1151) to about 20% of those in insulin-treated cells but did not increase pAKT or pERK levels (Fig. 5B). In the presence of insulin, S1-F8-S2 functioned as an antagonist, inhibiting insulin-dependent IR activation (Fig. S6A-E) and cell proliferation (Fig. S6F). These results are consistent with our structural observations (Fig. S3J) and indicate that S1-F8-S2 may disrupt the autoinhibitory conformation of IR, but does not stabilize IR in an active conformation.

**Figure 5.**
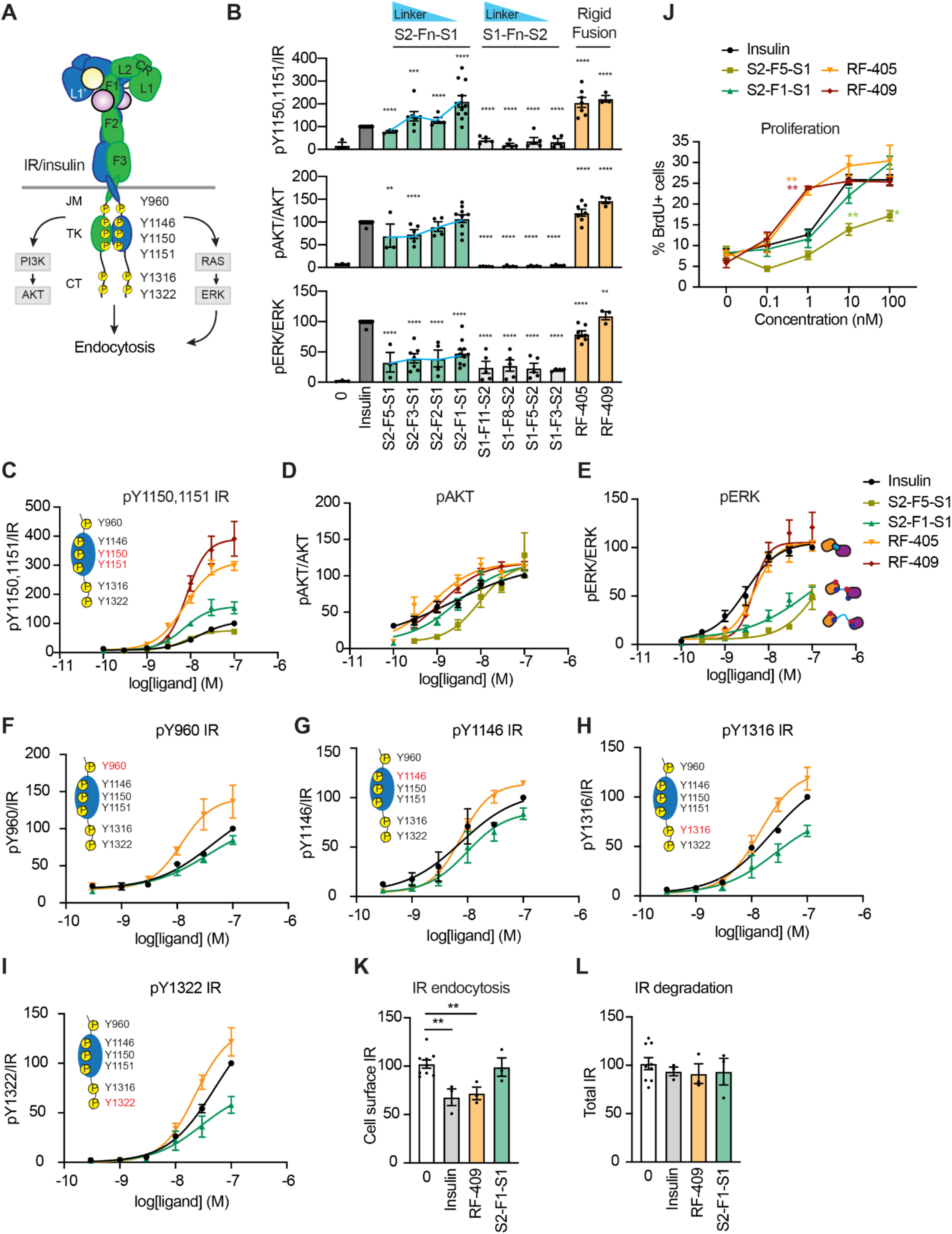
Varying binder orientation and stability results in different cell signaling outcomes. (A) Insulin triggers IR trans-autophosphorylation at multiple tyrosine residues in intracellular domains and activates two major signaling pathways. Activated IR undergoes endocytosis. The MAPK pathway promotes IR endocytosis. Key autophosphorylation residues in intracellular domains were shown. JM, juxtamembrane; TK, tyrosine kinase; CT, C-terminal domains of IR. (B) IR signaling in DKO-IR-B cells treated with the indicated ligands for 10 minutes. Mean ± sem. N = at least 3 independent experiments. Significance calculated using two-tailed student’s t-test. p values vs insulin. **p<0.01, ***p<0.001, and ****p<0.0001. (C-E) IR signaling in C2C12-IR cells by the indicated concentrations of ligands for 10 minutes. Levels of protein phosphorylation were normalized to total protein levels and shown as intensities relative to that in 100 nM insulin-treated cells. Data were fit by nonlinear regression. Mean ± sem. N= at least 3 independent experiments. (F-I) IR autophosphorylation in C2C12-IR cells by the indicated concentrations of ligands for 10 minutes. Levels of IR autophosphorylation were normalized to total IR levels and shown as intensities relative to that in 100 nM insulin-treated cells. Data were fit by nonlinear regression. Mean ± sem. N= at least 3 independent experiments. (J) Cell proliferation. C2C12-IR cells were incubated with the indicated concentrations of ligands for 24 hours and then incorporated Bromodeoxyuridine (BrdU) for 2 hours. BrdU-positive cells were analyzed by FACS. Mean ± sem. N= at least 3 independent experiments. Significance calculated using 2-way ANOVA. P value vs insulin. **p<0.01. (K) Quantification of cell surface IR at 30 minutes after incubation with 100 nM indicated ligands in primary mouse hepatocytes. Mean ± sem. N= 3 independent experiments. Significance calculated using two-tailed student’s t-test. p values vs insulin. **p<0.01. (L) Quantification of total IR at 30 minutes after incubation with 100 nM indicated ligands in primary mouse hepatocytes. Mean ± sem. N= 3 independent experiments.

In contrast to S1B-S2B fusion, we found that the flexibly-linked S2-Fn-S1 constructs functioned as partial (biased) agonists while the rigidly-linked RF-405 and RF-409 were full insulin mimicking, balanced agonists. The rigidly-linked constructs RF-405 and RF-409 elicited pY1150,1151 IR and pAKT and pERK levels similar to insulin, while S2-Fn-S1 constructs increased IR autophosphorylation (pY1150,1151 IR) and pAKT levels, but pERK levels were only 40% of those in insulin-treated cells (Fig. 5B; increasing the linker length reduces IR autophosphorylation). The increased dynamics of S2-Fn-S1 compared to the rigid RF-401 and RF-409 appears to compromise balanced signaling. We analyzed levels of pY1150,1151 IR, pAKT, and pERK over a wide range of ligand concentrations in C2C12-IR cells (Fig. 5C-E, S6G). RF-405 and RF-409 potently activated both pAKT and pERK levels similar to insulin, while the flexibly-linked S2-F1-S1 and S2-F5-S1 primarily activated pAKT (Fig. 5C-E, S6G), again indicating partial agonism. S2-F5-F1, which has a longer flexible linker, was significantly less effective at increasing pERK levels compared to S2-F1-S1 (Fig. 5E, S6G). A similar signaling pattern was observed with the single chain peptide S597^20,27,28^.

As S2-Fn-S1 is a partial agonist of IR signaling while RF-405 and RF-409 appear to be full balanced agonists, suggesting that altered dynamics (rather than structural changes per se) might lead to different IR autophosphorylation patterns. When insulin activates the IR, several tyrosine residues in the juxtamembrane (e.g. Y960), kinase (e.g. Y1146, Y1150, Y1151), and C-terminal domains (e.g. Y1316, Y1322) undergo trans-autophosphorylation (Fig. 5A). These phosphorylation sites are crucial for recruiting downstream substrates ^1,29^. To determine if the designed agonists differ in their efficiency at activating phosphorylation in these three domains, we measured the IR phosphorylation levels at these three intracellular sites. RF-405 increased autophosphorylation sites at all sites more significantly than insulin (Fig. 5F-I, S6G). While S2-F1-S1 increased pY1150,1151 levels more than insulin (Fig. 5C, S6G), phosphorylation at Tyr960 and Tyr1146 was lower (Fig. 5F,G, S6G), and in the C-terminal domain pY1316 and pY1322 phosphorylation was only 50% of insulin (Fig. 5H,I, S6G), indicating partial agonism. Thus conformational dynamics modulate the extent of IR phosphorylation during IR activation, which in turn modulates IR signaling.

The MAPK pathway regulates cell growth and proliferation^30^. We next investigated the differences in the extent of MAPK pathway (pERK levels) activation of our synthetic agonists on cell proliferation. We compared the ability of the designed agonists and insulin to induce proliferation of C2C12-IR cells (Fig. 5J). RF-405 and RF-409 induced cell proliferation at levels comparable to insulin. At 1 nM, cells treated with RF-405 and RF-409 showed a two-fold increase in proliferation compared to insulin. This enhanced proliferation may reflect the higher potency of RF-405 and RF-409 in inducing IR autophosphorylation. S2-F1-S1, which stimulated lower pERK levels, still induced cell proliferation, but S2-F5-S1 which is more defective in the MAPK pathway activation did not. Insulin-activated IR undergoes endocytosis which can terminate and redistribute IR signaling^31,32^. The MAPK pathway plays a crucial role in regulating IR endocytosis^33^ (Fig. 5A). To determine the effects of our synthetic agonists on IR endocytosis, we incubated S2-F1-S1, RF-409, or insulin with primary hepatocytes and analyzed cell surface IR and total IR levels. RF-409 and insulin, but not S2-F1-S1, significantly reduced cell surface IR levels (Fig. 5K; none of the three molecules reduced total IR levels, Fig. 5L). These data suggest that RF-409, like insulin, induces IR endocytosis, but S2-F1-S1 does not, supporting a role of the MAPK activation in promoting IR endocytosis.

Taken together, these data suggest that by modulating the relative orientation of the L1 and F1 domains of the IR during activation, different signaling and trafficking outcomes can be obtained. An equimolar mixture of unlinked S1B and S2B, did not induce IR phosphorylation or pERK or pAKT activation (Fig. S1H,I), indicating that engaging the two domains independently has no effect on IR activation. Combining them with rigid constructs to match and stabilize the active state of the IR results in insulin-like signaling and induction of endocytosis, whereas allowing a wide range of conformations with flexible linkers results in partial agonism (reduced pERK signaling) and less endocytosis.

### Designed IR agonists activate IR mutants that are resistant to insulins

Our fusion constructs and insulin make different sets of contacts with the IR: site-1 insulin interacts with the L1 domain and α-CT motif, while the S1B binds solely to the L1 domain (Fig. 6A). To test the importance of each binding site in activating IR, we first introduced mutations on the L1 domain of IR (F64A and F96A) that remove most of the interactions between L1 and insulin but only a subset of the interactions with the designed agonists (Fig. 6A; the designs have a much larger interaction surface area with L1 than insulin– 1481.8 vs 793.1 Å^2^ buried surface area respectively). As expected, insulin could not activate IR F64A and F96A (Fig. 6C,D), but the mutants could still be activated by our designed agonists. To evaluate the role of site-2 interface in activating IR, we introduced K484E and L552A mutations (Fig. 6B-D). Insulin and S2-F1-S1 could not activate the IR K484E/L552A mutants, supporting the importance of site2 interface on IR activation.

**Figure 6.**
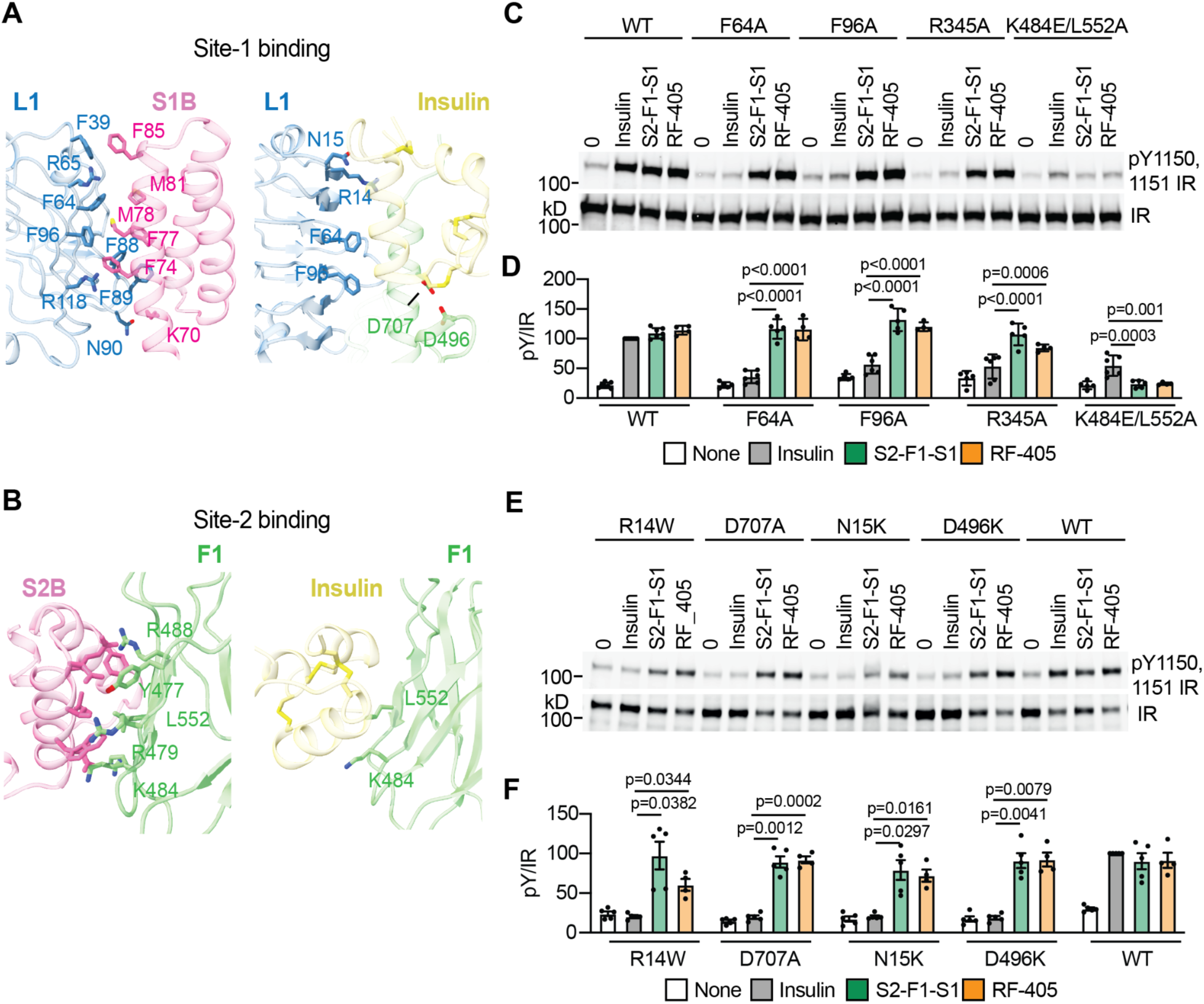
Designed IR agonists activate insulin resistant IR mutant. (A) Close up view of binding of RF-405 at IR L1 domain (left), and insulin at IR L1 domain (right). IR L1 domain is illustrated as transparent blue cartoons, while RF-405 and insulin are illustrated as pink and yellow cartoons, respectively. Major interacting residues are represented as sticks. (B) Close up view of binding of RF-405 at IR F1 domain (left), and insulin at IR F1 domain (right). IR F1 domain is illustrated as transparent green cartoons, while RF-405 and insulin are illustrated as pink and yellow cartoons, respectively. Major interacting residues are represented as sticks. (C) IR autophosphorylation by 10 nM insulin, S2-F1-S1, and RF-405 for 10 minutes in 293FT cells expressing WT IR or the indicated IR mutants. (D) Quantification of the western blot data shown in (C). Levels of IR autophosphorylation were normalized to total IR levels and shown as intensities relative to that in WT IR in insulin-treated cells. Mean ± SD. Significance calculated using 2-way ANOVA. N= at least 4 independent experiments. (E) IR autophosphorylation by 10 nM insulin, S2-F1-S1, and RF-405 for 10 minutes in 293FT cells expressing WT IR or the indicated disease-causing IR mutants. (F) Quantification of the western blot data shown in (E). Levels of IR autophosphorylation were normalized to total IR levels and shown as intensities relative to that in WT IR in insulin-treated cells. Mean ± sem. Significance calculated using Two-way ANOVA. N= at least 4 independent experiments.

Insulin induces a large conformational change and generates intra- and inter-domain contacts that stabilize the active IR state^8^. Arg345 in the L2 domain and Glu697 in the α-CT forms a salt bridge in the insulin-induced compact T-shaped IR and the R345A mutant is insulin resistant. We found that unlike insulin, S2-F1-S1 and RF-405, were able to fully activate the IR R345A mutants (Fig. 6C,D). Therefore, the activation of IR by designed agonists is less dependent on this Arg-Glu salt bridge formation and the binding energy of the designed agonists is sufficient to overcome this loss of internal stabilization of the active state.

Mutations in the insulin binding sites of IR cause rare but severe insulin resistance syndromes such as Donohue syndrome and Rabson-Mendenhall syndrome^34–36^. Patients produce insulin normally, but are not able to properly regulate glucose metabolism. We introduced disease-causing mutations in the L1 (R14W^37^ and N15K^38^), F1 (D496N^39^, D496K) and α-CT (D707A) ^40^ domains of IR at sites that do not contribute to the interaction with the designed IR agonists (Fig. 6E,F). As expected, insulin could not activate the R14W, N15K, D496K, and D707A IR mutants. In contrast, both S2-F1-S1 and RF-405 could fully activate these disease-causing mutants (Fig 6E,F). These results highlight the differences in activation mechanism of our designed agonists, and such designed agonists could be beneficial for patients with insulin-binding deficient IR mutants.

### Designed agonists mimic insulin functions *in vivo*

We set out to investigate the function of our designed agonists *in vivo*. The IR structure and sequence are highly conserved between humans and mice. We first tested if the designed agonists can activate IR in mouse cells (Fig. S7A,B). We isolated primary mouse hepatocytes and compared insulin- and designed agonist-induced IR signaling. In primary hepatocytes, RF-409 increased IR autophosphorylation (pY1150,1151 IR) and pAKT similarly to insulin, while S2-F1-S1 induced pY1150,1151 IR and pAKT less potently than RF-409. Due to the high basal levels of pERK in primary hepatocytes, despite insulin stimulation, we could not observe significant increase in pERK. We next compared IR signaling induced by S2-F1-S1 and RF-409 in metabolic tissues including liver and skeletal muscle (Fig. S7C,D). In both tissues, S2-F1-S1 was less effective in stimulating IR autophosphorylation, but it could increase levels of pAKT to similar levels as RF-409. As in primary hepatocytes, mouse liver exhibited high basal levels of pERK, and we did not observe significant difference between all tested molecules. In skeletal muscle, RF-409 significantly increased pERK levels, while S2-F1-S1 did not, confirming partial agonism *in vivo*.

To determine the metabolic effects of our designed agonists, we conducted an insulin tolerance test (ITT) in mice fed with a normal diet (Fig. 7A,B; Fig. S7E,F). S2-F1-S1 reduced glucose levels in mice as effectively as insulin, while RF-409 was even more effective than insulin. The designed agonists had longer-lasting effects on glucose levels compared to insulin: with insulin, glucose rapidly dropped (within 30 minutes) and began to rise again shortly thereafter, while with half the dose of RF-409, glucose levels decreased to the same level and then remained low throughout the experiment. To determine the physiological effects of designed agonists under diabetic conditions, we conducted ITT in diet-induced obese mice, which have higher basal blood glucose levels. Similar to healthy mice, RF-409 and S2-F1-S1 slowly reduced glucose levels but exhibited prolonged glucose lowering effects compared to insulin. RF-409 was more effective at lowering glucose levels than S2-F1-S1 (Fig. 7C; Fig. S7G). Strikingly, after a single injection of RF-409, mice maintained low glucose levels for 6 h, whereas mice treated with insulin rose glucose levels within 2 h (Fig. 7D; Fig. S7H). Overall, our designed agonists effectively lower glucose levels in mice, with RF-409 proving to be more potent than S2-F1-S1 (Fig. 7B,C; Fig. S7F).

**Figure 7.**
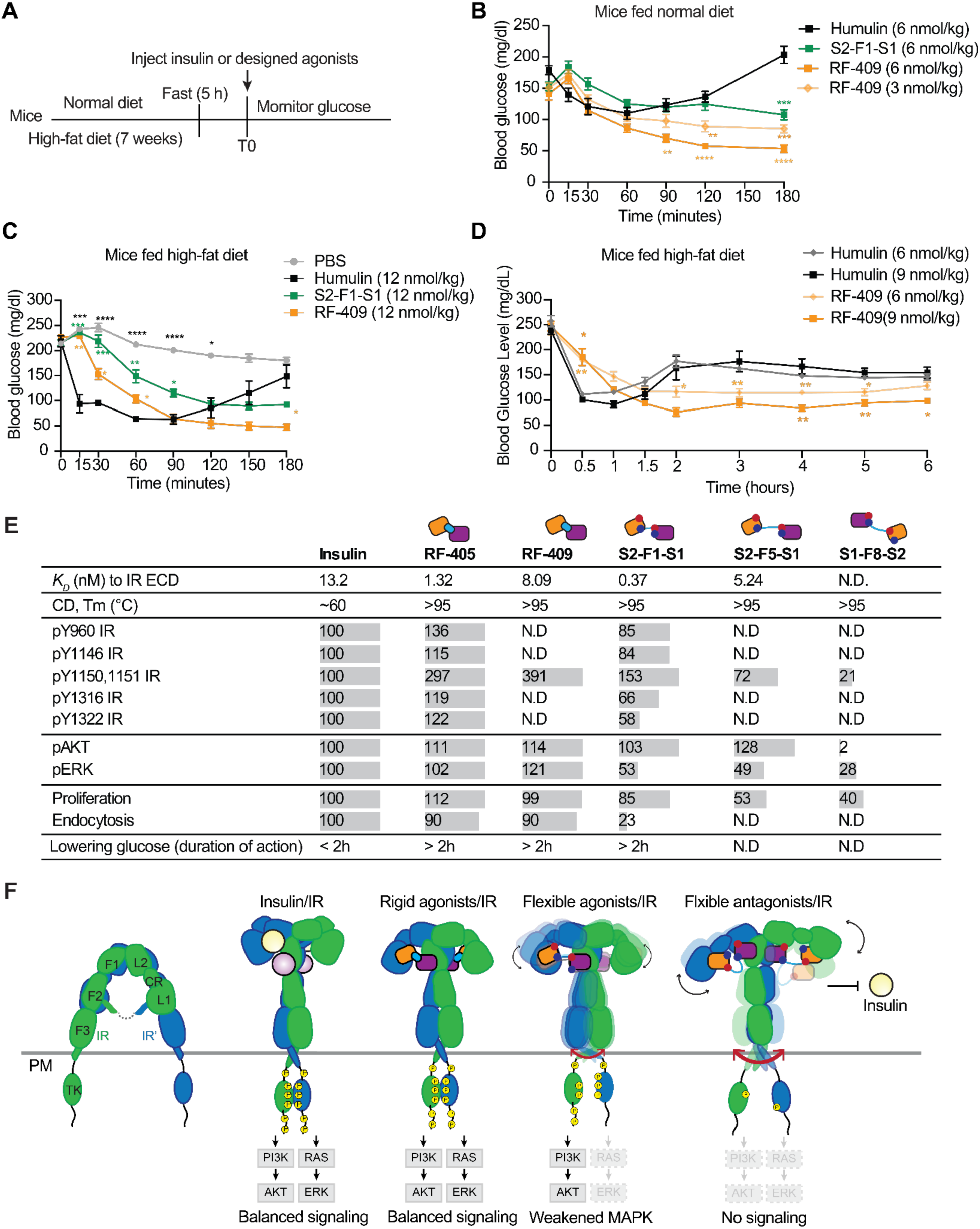
Designed IR agonists tune IR signaling and reduce glucose levels. (A) Illustration of the mouse experiment with normal chow and high-fat diets. 2- to 3-month-old male mice were used. (B) Insulin tolerance test in mice fed normal chow diet. Mice were injected intraperitoneally with Humulin, S2-F1-S1 or RF-409 at the indicated doses, and their blood glucose levels measured at the indicated time points after injection. Mean ± sem. Humulin, N=8; S2-F1-S1 and RF-409, N=6. Significance calculated using 2-way ANOVA. p value vs Humulin. **p<0.01, ***p<0.001, and ****p<0.0001. (C) Insulin tolerance test in mice fed a high-fat diet. Mean ± sem. Humulin, N=8; S2-F1-S1 and RF-409, N=7 mice per group. Significance calculated using 2-way ANOVA. p value vs Humulin. *p<0.05, **p<0.01, ***p<0.001, and ****p<0.0001. (D) Insulin tolerance test in mice fed a high-fat diet. Mean ± sem. Humulin, N=8; S2-F1-S1 and RF-409, N=7 mice per group. Significance calculated using 2-way ANOVA. p value vs Humulin with the same dose. *p<0.05, **p<0.01, ***p<0.001, and ****p<0.0001. (E) Overview of efficacy (Emax relative to insulin-activated IR WT) and property of IR agonists. Mean ± SD. Significance calculated using 1-way ANOVA, p value vs insulin. *p<0.05, **p<0.01, ***p<0.001, and ****p<0.0001. Bar was shown to represent fold changes compared to insulin functions. Values greater than insulin were displayed in the same size bar as insulin maximum values. N.D, not determined. (F) IR structural stability during activation is essential for downstream signaling, trafficking, and biological function.

It has long been thought that IR-mediated AKT pathway controls metabolic function, while the MAPK pathway controls mitogenic function. However, molecular dissection of the two pathways has been hindered by a major hurdle: the native ligand of IR, insulin activates both pathways. Our series of designed agonists provide insight into the mechanism of signaling through the IR, and how receptor autophosphorylation is tied to downstream outcomes. RF-405, S2-F1-S1 and S2-F5-S1 all bind to the IR with high affinity (Fig. 3, 7E) but generate different signal transduction processes. The rigid RF-405 agonist induces a highly ordered active conformation of IR (Fig. 4) which leads to efficient IR autophosphorylation in the juxtamembrane (pY960), tyrosine kinase (pY1150,py1151) and C-terminal (pY1316 and pY1322) domains. This results in strong activation of the PI3K-AKT (pAKT), and MAPK (pERK) pathways (Fig. 5C-I, 7E, Fig. S6G). The intermediate flexibility agonist S2-F1-S1 induces a more dynamic conformation of the receptor that is less able to autophosphorylate, particularly the C-terminal domain (Y1316 and Y1322), which reduces activation of the MAPK pathway and endocytosis. SHP2^41^, an upstream activator of the MAPK pathway, docks on the IR following phosphorylation at the C-terminal tyrosine sites, and modulates IR signaling by dephosphorylating IR and IR substrates and promoting receptor endocytosis ^31,33,42–45^. S2-F1-S1 is likely less effective in activating the MAPK pathway and IR endocytosis because it does not induce phosphorylation at these sites. S2-F1-S1 was less effective in controlling glucose levels in mice than the rigid agonists, suggesting an important role of the MAPK pathway in regulating metabolism (S2-F1-S1 does induce the AKT pathway). The highly flexible S2-F5-S1 showed even weaker MAPK pathway activation (Fig 5B, 5E, 7E), while S1-F8-S2 does not favor an active conformation and hence does not signal, but does disrupt the autoinhibitory state of IR and is functionally antagonist, competing out insulin.

### Limitations

Although our flexible agonists showed biased agonism in cultured cells, we have not demonstrated *in vivo* effects of the biased agonism on glucose/lipid metabolism and tissue homeostasis. The relatively large size of designed agonists compared with native insulin could limit penetration of skeletal muscle and adipose tissue, without affecting delivery to the liver. The slow yet prolonged glucose lowering effects of designed agonists could reflect their tissue selectivity *in vivo*: further investigation will be necessary to explore the pharmacokinetics and long-term efficacy of designed binders *in vivo*.

## Discussion

Understanding how physiological signals are transmitted to downstream pathways, particularly those involving conformational dynamics, has been challenging. Here, we demonstrate that computationally designed agonists with controlled extents of internal flexibility induce states of the target receptor with matching dynamics. We describe rigid and flexible agonists that bring together the receptor site-1 and site-2 containing domains and drive the receptor into a T-shaped active state, but with different dynamics as indicated by the cryo-EM structure resolution. Comparison of the effects of the rigid and flexible agonist shows that the extent of ordering directly impacts autophosphorylation of IR during activation, MAPK versus AKT downstream signaling, and trafficking, indicating that the conformational stability of IR during activation is a key factor in controlling its biological activity (Fig. 7F). The designed agonists are also highly stable and easy to produce recombinantly, making them ideal candidates for large-scale production and storage, and to our knowledge are the most potent completely synthetic IR agonists available described to date.

Moving forward, there are a number of exciting avenues to explore. First, our agonist design approach enables tuning the conformation and dynamics of the active IR, which cannot be readily explored with native insulin. By varying the geometry of rigid fusions using RFdiffusion, and the linker length and composition in the flexible fusions, it should be possible to modulate the pattern of IR autophosphorylation, MAPK signaling intensity, intracellular IR trafficking, and cell proliferation. Given the potential cancer inducing properties of insulin ^46–49^, such partial agonists could be beneficial for patients with both diabetes and cancer. Second, our designed agonists exhibit high binding affinity and selectivity for IR, and it will be useful to explore whether this is useful under conditions where off-target activation of IGF1R is problematic such as cancer, autoimmune disease, and thyroid eye disease ^50–52^. Third, as our designed agonists bind and activate insulin-binding deficient and disease-causing IR mutants, they could provide the basis for therapeutic interventions for these rare but devastating conditions that currently lead to early morbidit

## STAR Methods

### Computational design of the IR site 2 binders

The computational design method was using the method previously described^17^. In brief, IR structure 6PXV was downloaded from Protein Data Bank and relaxed by Rosetta guided by experimental design-guided relaxation. The F1 domain (residue 467-590) was extracted as the targeting domain and the insulin binding side was selected as the targeting interface. For each residue of the selected interface, Rotamer Interaction Field (RIF) was generated. Later, mini-protein scaffold set, composed of 3 helical, 4 helical and ferredoxin scaffolds, were used to search for global shape complementarity with PatchDock. The docked scaffolds were then sequenced sequence-optimized using Rosetta FastDesign and evaluated by DeepAccNet^18^ pLDDT and Rosetta Metrics including ddG, contact patch, and contact molecular surface^17^. A total of 11280 oligos encoding designed site 2 binders passed the filters.

### Computational design of the linked binders

Flexibly-linked IR binders were generated by linking S1B and S2B with Gly-Ser (GS) linkers with various lengths. Non Interface residues 1, 13, 22, 25, 27, 29, 30, 35, 36, 39, 43, 44, 45, 48, 51, 52, 53, 54, 55, 58, 59, 60, 62, 64 were redesigned by MPNN to increase the solubility of S2-F1-S1.

To generate Rigidly-fused IR binders, we aligned the design models of S1B/L1 and S2B/F1 to the S2-F1-S1/IR complex as the starting point. The first helix of S1B (Up to residue 3) and the last helix of S2B (Residue 54 onwards) were rebuilt by RFdiffusion to link the L1- and F1-binding interfaces. Surface residues near the diffused regions were masked and redesigned using ProteinMPNN along with the newly built regions. 32 sequences were generated for each backbone. Then, the designs were predicted by AlphaFold 2 (AF2), relaxed and scored by Rosetta. The top 28 designs with the highest AF2 pLDDT and lowest RMSD to design were selected for experimental characterization. Sequence optimization was further performed to improve solubility of the Rigidly-linked IR binders.

### Yeast surface display screening for IR site 2 binders with FACS

The yeast surface display screening was performed using the protocol as previously described^14,16^. Briefly, DNAs encoding the minbinder sequences were transformed into EBY-100 yeast strains. The yeast cells were grown in CTUG medium and induced in SGCAA medium. After washing with FACS-buffer (PBS, Fisher Scientific, supplemented with 1% w/v bovine serum albumin, SigmaAldrich), the cells were incubated with 1uM biotinylated IR F1 domain together with streptavidin–phycoerythrin (SAPE, ThermoFisher, 1:100) and anti-c-Myc fluorescein isothiocyanate (FITC, Miltenyi Biotech, 6.8:100) for 60 min. After washing twice with FACS buffer, the yeast cells were then resuspended in the buffer and screened via FACS. Only cells with PE and FITC double-positive signals were sorted for next-round screening. After another round of enrichment, the cells were titrated with biotinylated IR F1 domain at 100 nM, 10 nM and 1 nM for 60 min, washed, and further stained with both streptavidin–phycoerythrin (SAPE, ThermoFisher) and anti-c-Myc fluorescein isothiocyanate (FITC, Miltenyi Biotech) at 1:100 ratio for 30 min. After washing twice with FACS buffer, the yeast cells at different concentrations were sorted individually via FACS and regrown for 2 days. Next, the cells from each subpool were lysated and their sequences were determined by next-generation sequencing.

### Protein binder expression and purification

Synthetic genes encoding designed proteins were purchased from Genscript or Integrated DNA Technologies (IDT) in the pET29b expression vector or as eBlocks (IDT) and cloned into customized expression vectors^53^ using Golden Gate cloning. A His6x tag was included either at the N-terminus or the C-terminus as part of the expression vector. Proteins were expressed using autoinducing TBII media (Mpbio) supplemented with 50x5052 and 20 mM MgSO_4_ in BL21 DE3 *E.coli* cells. Proteins were expressed under antibiotic selection at 25 °C overnight after initial growth for 6-8 h at 37 °C. Cells were harvested by centrifugation at 4000x g and resuspended in lysis buffer (20 mM Tris, 300 mM NaCl, 5 mM imidazole, pH 8.0) containing protease inhibitors (Thermo Scientific) and Bovine pancreas DNaseI (Sigma-Aldrich) before lysis by sonication.

Proteins were purified by Immobilized Metal Affinity Chromatography (IMAC). Cleared lysates were incubated with 0.1-0.5 mL nickel NTA beads (Qiagen) for 20-40 minutes before washing beads with 5-10 column volumes of lysis buffer, 5-10 column volumes of wash buffer (20 mM Tris, 300 mM NaCl, 30 mM imidazole, pH 8.0). Proteins were eluted with 1-4 mL of elution buffer (20 mM Tris, 300 mM NaCl, 300 mM imidazole, pH 8.0). All protein preparations were as a final step polished using size exclusion chromatography (SEC) on Superdex 75 Increase 10/300GL columns (Cytiva) using PBS buffer (Fisher Scientific). SDS-PAGE and LC/MS were used to verify peak fractions. Proteins were concentrated to concentrations between 0.5-10 mg/mL and stored at room temperature or flash frozen in liquid nitrogen for storage at -80. Thawing of flash-frozen aliquots was done at room temperature. All purification steps from IMAC were performed at ambient room temperature.

### Biolayer interferometry (BLI)

The BLI experiments were performed on an OctetRED96 BLI system (ForteBio) at room temperature in HBS-EP buffer (Cytiva Life Sciences) supplemented with 0.2 % w/v bovine serum albumin (BSA, SigmaAldrich). Prior to measurements, streptavidin-coated biosensors were first equilibrated for at least 10 min in the assay buffer. Biotinylated target proteins (IR F1 domain, IR extracellular domain (ECD) (ACROBIOsystems INR-H82E6), IGF1R ECD (ACROBIOsystems IGR-H82E3)) were immobilized onto the biosensors by dipping them into a solution with 100 nM protein until the loading signal reaches 0.5 - 1 nm. Association and dissociation of the analytes were monitored by dipping biosensors in the solutions containing analytes at various concentrations for 300s followed by dipping the biosensors in fresh buffer for 300s. Experiments were performed at 25 °C while rotating at 1000 rpm. Global kinetic or steady-state fits were performed on buffer-subtracted data using the manufacturer’s software (Data Analysis 12.1) assuming a 1:1 binding model.

### Circular Dichroism Spectroscopy

CD spectra were recorded in a 1 mm path length cuvette at a protein concentration between 0.3-0.5 mg/mL on a J-1500 instrument (Jasco). For temperature melts, data were recorded at 222 nm between 25 and 95 °C every 2 °C, and wavelength scans between 190 and 260 nm at 10 °C intervals starting from 25 °C. Experiments were performed in 10 mM sodium phosphate buffer (pH 7.4), 50 mM NaCl. The high tension (HT) voltage was monitored according to the manufacturer’s recommendation to ensure optimal signal-to-noise ratio for the wavelengths of interest.

### Protein expression and purification for cryo-EM

For structural studies, the short isoform of human insulin receptor (hIR) or mouse insulin receptor (mIR, sharing 94% sequence homology with hIR) were cloned into pEZT-BM expression vectors as described previously^8–10^. To improve expression and protein behavior, seven mutations (Y960F, S962A, D1120N, R1333A, I1334A, L1335A, L1337A: amino acid numbering of short isoform of hIR without signal peptide) were introduced to hIR, and two mutations (Y962F and D1122N, amino acid numbering of short isoform of mIR without signal peptide) were introduced to mIR. The Human Rhinovirus 3 C recognition site (3C), affinity purification tag Tsi3 (T6SS secreted immunity protein three from *Pseudomonas aeruginosa*) and His_8_ tag were fused to the C-terminus of both proteins.

The expression and purification of hIR and mIR were performed following previously described protocols with minor modifications^8–10^. Briefly, the plasmids were transformed to *Escherichia coli* strain DH10Bac to produce bacmid DNA. Recombinant baculovirus was generated by transfecting Sf9 cells with bacmid DNA using Cellfectin reagent (Gibco). hIR or mIR proteins were expressed in FreeStyle 293-F cells by infecting the cells with the virus at 1:10 (virus: cell, v/v) ratio. Six hours after infection, 8 mM sodium butyrate was added to boost protein expression. Cells were cultured in a shaking incubator supplemented with 8% CO_2_ for 48-60 h at 30 °C before harvesting.

The cells were resuspended in lysis buffer containing 40 mM Tris-HCl pH 7.5, 400 mM NaCl (Buffer A) with Protease Inhibitor Cocktail (Roche) and lysed by using a French Press cell disruptor. The membrane fraction was obtained by ultracentrifugation of the cell lysate for 1 h at 100,000 g at 4 °C. To extract the protein from the membrane fraction, Dodecyl maltoside (DDM, Anatrace) was added to a final concentration of 1% (m/v) with stirring overnight. The supernatant containing the solubilized protein was obtained by ultracentrifugation for 1 h at 100,000 g at 4 °C. The supernatant was added with 2 mM CaCl_2_ and Tse3 protein-conjugated Sepharose resin (GE Healthcare) and incubated at 4 °C for 1 h before being loaded onto a column by gravity flow. The resin was subsequently washed with 40 column volumes (CV) of buffer containing 40 mM Tris-HCl pH 7.5, 400 mM NaCl, 2 mM CaCl2, 5% glycerol (v/v), 0.05% DDM (m/v) (Buffer B) and eluted by HRV-3C protease cleavage at 4 °C overnight. The protein was then concentrated using a 100 kDa cutoff concentrator (Millipore), loaded onto a Superose 6 increase 10/300 GL size-exclusion column (Cytiva), and eluted with buffer containing 20 mM HEPES pH 7.4, 150 mM NaCl and 0.03% DDM (Buffer C). The dimer fractions of mIR or hIR proteins were identified by SDS-PAGE and pooled.

To make Insulin/S2B/mIR complex for cryo-EM analyses, commercial insulin (I2643, recombinantly expressed in yeast) and site-2 binder (S2B) were added to mIR at a molar ratio of 4:4:1 (insulin: S2B: mIR). To make RF-405/hIR, S2-F1-S1/hIR, and S1-F8-S2/hIR complexes, RF-405, S2-F1-S1, S1-F8-S2 binders were added to hIR at a molar ratio of 4:1 (binder: hIR). After incubation for 30 min, the protein mixtures were concentrated to 6-8 mg ml^-1^ using 100 kDa cutoff concentrators (Millipore) and subject to cryo-EM grid preparation immediately. All purification and following steps were performed at 4 °C or on ice.

For analytical size-exclusion chromatography of S2-F1-S1/hIR and RF-405/hIR complexes, S2-F1-S1 or RF-405 were added to hIR at a molar ratio of 4:1 (binder: hIR). After incubation for 30 min, the protein mixtures were then loaded onto a Superose 6 increase 3.2/300 analytical size-exclusion column (Cytiva) and eluted with Buffer C.

### Cryo-EM data collection and image processing

EM data acquisition, image processing, and model building, and refinement were performed following previous protocols with some modifications^8–10,20^. The samples of IR in complex with insulin/S2B, RF-405, or S2-F1-S1 were applied to glow-discharged Quantifoil R1.2/1.3 300-mesh gold holey carbon grids (Quantifoil, Micro Tools GmbH, Germany). Grids were blotted under 100% humidity at 4 °C and plung-frozen in liquid ethane using a Mark IV Vitrobot (Thermo Fisher Scientific). Micrographs were collected in the counting mode on either Glacios or Titan Krios microscopes (Thermo Fisher Scientific) with either Falcon4 (Thermo Fisher Scientific) or K3 Summit direct electron detectors (Gatan). The nominal magnification and pixel size of each data set are summarized in Tables S1.

Motion-correction and dose-weighting of the micrographs were carried out using the Motioncor2 program (version 1.2)^54^. GCTF 1.06 was used for CTF correction^55^. Template-based particle picking was carried out using the autopick tool in RELION 4.0^55,56^. Particles were cleaned up with multiple rounds of 2D and 3D classification in RELION. Good particles were selected and subjected to 3D refinement with C2 symmetry. The exact procedures are summarized in supplementary figures. The initial mode for 3D classification and refinement was generated using the SGD method in RELION. The refined maps were further improved by using Bayesian polishing and CTF refinement at the final stage. The Fourier Shell Correlation (FSC) 0.143 criterion was used for estimating the resolution of the maps. Local resolution was calculated in RELION.

### Model building and refinement

To build the atomic models of the IR structures with different binders bound, the published model of each domain of human IR (PDB ID: 6PXV) and the predicted models of S2B, RF-405 or S2-F1-S1 using AlphaFold2 ^57^ were docked into the cryo-EM maps as rigid-body. The models were adjusted manually in Coot 0.98 ^58^. The models were refined using the real-space refinement module in Phenix 1.18 ^59^. Model quality was checked using Molprobity as a part of the Phenix validation tool set ^60^. Model statistics are summarized in Table S1. Structural figures were rendered in ChimeraX 1.7 ^61,62^.

### Mouse strains and Husbandry

Animal work described in this manuscript has been approved and conducted under the oversight of the Columbia University Institutional Animal Care and Use Committee. Mice (C57BL/6J, Jackson Laboratory, #000664) were fed a standard rodent chow (Lab diet, #5053) or high-fat diet (HFD) (D12492; Research Diets). All animals were maintained in a specific antigen-free barrier facility (temperature, 20-26 °C; humidity, 30-70%) with 12 h light/dark cycles (6 a.m. on and 6 p.m. off). Two to three-month-old male mice were used in this study.

### Cell cultures, Transfection, and Viral Infection

IR and IGF1R double knockout brown preadipocytes expressing only human IR-B (DKO-IR-B) or mouse IR-A (DKO-IR-A) were kindly provided by Dr. Ronald Kahn^63^. DKO-IR-B, DKO-IR-A, 293FT (Invitrogen, #R70007), and C2C12 (ATCC, CRL-1722) were cultured in high-glucose DMEM supplemented with 10% (v/v) FBS, 2 mM L-glutamine, and 1% penicillin/streptomycin and maintained in monolayer culture at 37 °C and 5% CO2 incubator.

Plasmid transfections into 293FT cells were performed with LipofectamineTM 2000 (Invitrogen). To generate C2C12 cells expressing human IR-A, 293FT cells were transfected with pBabe-IR-A-GFP, pCMV-gag/pol, and pCMV-VSV-G. Virus were collected at 2- and 3-days after transfection and concentrated with homemade virus concentrator. C2C12 cells were infected with concentrated virus and polybrene (4 ug/ml). Cells were selected with 2 mg/ml of puromycin at 3 days after infection and sorted using FACS sorter (Sony Ma900). To generate IGF1R knockout C2C12 cells expressing human IR-A (C2C12-IR), lentiviruses were packaged in 293FT cells by transfecting the cells with lentiCRISPR vector (Addgene # 52962, gRNA: CACCGCTATGGTGGAGAGGTAACAG), psPAX2 (Addgene #12260), and pMD2.G (Addgene #12259). Viruses were collected at 2- and 3-days after transfection and concentrated. C2C12 cells expressing IR-A were infected with concentrated virus and polybrene (4 mg/ml). Cells were selected with blasticidin (10 mg/ml) for 3 weeks. Following passage of an aliquot of each cell line for three to four weeks, a fresh batch of cells was thawed and propagated. There were no signs of mycoplasma contamination.

### IR signaling assay

The IR signaling assay was performed as described earlier with some modification^9,10,20,64^. For IR mutants assay, 293FT cells were transfected with Myc-tagged IR mutants or WT. One day later, the cells were serum starved for 14-16 h. Serum-starved cells were treated with insulin (I2526, Sigma) or designed binders for 10 min. For binder validation, DKO-IR-A, DKO-IR-B, or C2C12-IR cells were used. Two days after seeding, the cells were serum starved for 6 h. Serum-starved cells were treated with insulin or designed binders for 10 min. To analyze antagonistic effects of binders, cells were serum starved for 4 h, treated with the indicated concentrations of binders for 1 h, and then treated for 10 min with insulin at the indicated concentrations.

After treatment, cells were incubated with cell lysis buffer B [50 mM Hepes pH 7.4, 150 mM NaCl, 10% (v/v) Glycerol, 1% (v/v) Triton X-100, 1 mM EDTA, 10 mM sodium fluoride, 2 mM sodium orthovanadate, 10 mM sodium pyrophosphate, 0.5 mM dithiothreitol (DTT), 2 mM phenylmethylsulfonyl fluoride (PMSF)] supplemented with cOmplete Protease Inhibitor Cocktail (Roche), PhosSTOP (Roche), and 25 U/ml turbo nuclease (Accelagen) on ice for 1 h. After centrifugation at 18,213 g at 4°C for 20 min, cell lysate samples were made with SDS-PAGE protein loading buffer. Cell lysates were analyzed by SDS-PAGE and Western blotting. Anti-IR-pY1150/1151 (1:2000, 19H7, Cell signaling; labeled as pY IR (or pY IGF1R), #3024), anti-IR-pY1146 (1:1000, D6D5L, Cell signaling, #80732S), anti-IR-pY960 (1:1000, Invitrogen, #44-800G), anti-IR-Y1316 (1:1000, Invitrogen, #44-807G), anti-IR-Y1322 (1:1000, Invitrogen, #44-809G), anti-Myc (1:2000; 9E10, Roche; labeled as IR, #11667149001), anti-IR (1:500; CT3, Santa Cruz, sc-57342), anti-AKT (WB, 1:2000; 40D4, #2920), anti-pS473 AKT (WB, 1:2000; D9E, #4060), anti-ERK1/2 (WB, 1:2000; L34F12, #4696), and anti-pERK1/2 (WB, 1:2000; 197G2, #4377) were used as primary antibodies. For quantitative Western blots, anti-rabbit immunoglobulin G (IgG) (H+L) (Dylight 800 conjugates, #5151) and anti-mouse IgG (H+L) (Dylight 680 conjugates, #5470) (Cell signaling) were used as secondary antibodies. The membranes were scanned with the Odyssey Infrared Imaging System (LI-COR, Lincoln, NE).

### IR signaling analysis *in vivo*

IR signaling *in vivo* analysis was performed as described earlier with some modifications^10,20,65,66^. 2-3-months-old male mice were fasted overnight. Following anesthesia, mice were injected with 6 nmol Humulin (Eli Lilly) or 9 nmol designed agonists per mouse via inferior vena cava. Livers and skeletal muscle were removed at 5 min and 10 min after injection, respectively. Tissues were homogenized in cell lysis buffer B supplemented with cOmplete Protease Inhibitor Cocktail (Roche), PhosSTOP (Sigma), and 25 U/ml turbo nuclease (Accelagen), homogenized with FisherbrandTM Bead Mill homogenizer, and then incubated on ice for 1hr. After centrifuge at 20,817 g at 4°C for 30 min, the concentrations of cell lysate were measured using Micro BCA Protein Assay Kit (Thermo Scientific). The lysates were then analyzed by quantitative western blotting (Li-COR, Lincoln, NE).

### Primary mouse hepatocytes isolation

Mouse primary hepatocytes were isolated from 2-3-month-old male mice with a standard two-step collagenase perfusion procedure as described earlier^20,66^. Isolated hepatocytes were resuspended with attached medium [Williams’ Medium E supplemented with 5% (v/v) FBS, 10 nM insulin, 10 nM dexamethasone, and 1% penicillin/streptomycin] and plated on collagen (Sigma, C3867)-coated dishes. After 4 h, the medium was changed to serum free low-glucose DMEM supplemented with 1% penicillin/streptomycin. After 14-16 h, the cells were treated with insulin to analyze IR signaling and IR endocytosis.

### Cell surface biotinylation and streptavidin pulldown

Cell Surface labeling was performed as previously described with some modifications^67^. Primary mouse hepatocytes were treated with 100 nM ligands for 30 min. Cells were washed with in cold PBS, pH 8.0 (Corning, 21-030-CM) and incubated in 0.5 mg/ml EZ-link^TM^ Sulfo-NHS-LC-Biotin (Thermo Fisher, 21335) dissolved in PBS, pH 8.0 on ice for 10 min. The labeling reaction was quenched in 50 mM glycine in two sequential 10 min incubations on ice. Cells were incubated with RIPA buffer (Thermo Fisher, 89901) supplemented with Halt Protease & Phosphatase inhibitor cocktail (Thermo Fisher, 78442) on ice for 1 h. After centrifugation at 18,213 g at 4°C for 20 min, the supernatant was taken for streptavidin pulldown. Lysates were incubated with streptavidin magnetic beads (Thermo Fisher, 88817) overnight and washed 3 times with RIPA buffer. Beads were eluted with SDS sample buffers and samples were analyzed by SDS-PAGE and Western blotting. Anti-IR (1:500; CT3, Santa Cruz, sc-57342), anti-beta catenin (1:1000, D10A8, Cell signaling, #8480), anti-Actin (1:1000, C4, Santa Cruz, #sc-47778).

### Insulin tolerance test

Mice were fasted for 2h (healthy mice) or 5 h (diet-induced obese mice) and their blood glucose levels (T=0) were measured with tail bleeding (Contour Next). Mice were then injected intraperitoneally with PBS, Humulin or designed agonists. Their blood glucose levels at the indicated time points after injection were measured with tail bleeding.

### Cell proliferation assay

C2C12-IR cells were seeded in 0.1 million cells per 35 mm cell culture dish. One day later, cells were serum starved for 24 h and then treated for 24 h with indicated concentrations of binders in the presence or absence of insulin. One day later, cells were pulsed with 10 μM bromodeoxyuridine (BrdU) for 2 h, then fixed with 70% cold ethanol. The fixed cells were washed with FACS blocking solution (0.02% Triton X-100 and 1% BSA in PBS), denatured with 3N HCl for 30 min, neutralized with phosphate/citrate buffer for 10 min, and then washed three times with the FACS blocking solution. 10uL of FITC-anti-BrdU antibody (BD, # 556028) was added to each sample and incubated at RT for 2.5 h. cells were washed with the FACS blocking solution and stained with propidium iodide (BD, #550825). Cells were analyzed using BD FACSCanto II from the Flow Cytometry Core of the Columbia Center for Translational Immunology (CCTI) and Herbert Irving Comprehensive Cancer Center (HICCC). Data was processed with FlowJo.

### Data statistical analysis

Prism 10 was used for the generation of graphs and for statistical analyses. Results are presented as mean ± s.d. or mean ± s.e.m. Two-tailed unpaired t tests were used for pairwise significance analysis. For dose-response curve, the relative intensity signals were fitted to a four-parameter sigmoidal concentration-response curve, from which the pEC_50_ values (negative logarithmic values of half-maximum effective concentration (EC_50_) values) were used to calculate the mean and s.e.m. Log-transformed EC50 values were analyzed by Extra sum-of-squares F Test in Prism, between insulin and designed agonists. Two-way ANOVA followed by Tukey’s multiple comparisons tests were used in case of more than two factors’ comparisons. No power analysis for sample sizes were performed. Randomization and blinding methods were not used, and data were analyzed after the completion of all data collection in each experiment.

## Author contributions

X.W., E.C., X.B., and D.B. designed the research. X.W and P.V. designed and characterized the biochemical properties of designed binders. S.C., M.N., C.H., and E.C. characterized the designed binders in cultured cells and analyzed cellular assays. A.H., S.C., M.H., D.A., and E.C. characterized the designed binders in mice and analyzed animal experiments. K.C., J.L., W.A., and X.C prepared samples, collected data, and built the cryo-EM models. X.W., E.C., X.B., and D.B. wrote the initial manuscript. All authors contributed to the edition and discussion of the manuscript.

