## Supplemental for "Tuning Insulin Receptor Signaling Using *De Novo* Designed Agonists"

Figure S1. Effects of site-1 binder and site-2 binder. Related to Fig2.
Figure S2. Cryo-EM analysis of the Insulin/S2B/IR complex. Related to Figure. 2.
Figure S3. Structures of RF-405/IR, S2-F1-S1/IR, and S1-F8-S2/IR. Related to Figure 4.
Figure S4. Cryo-EM analysis of the RF-405/IR complex. Related to Figure. 4.
Figure S5. Cryo-EM analysis of the S2-F1-S1/IR complex. Related to Figure. 4.
Figure S6. Antagonistic effects of S1-F8-S2. Related to Figure 5.
Figure S7. Designed IR agonists activate IR signaling and control glucose levels in mice. Related to Figure 7.
Table S1. Cryo-EM data collection and structure refinement statistics.

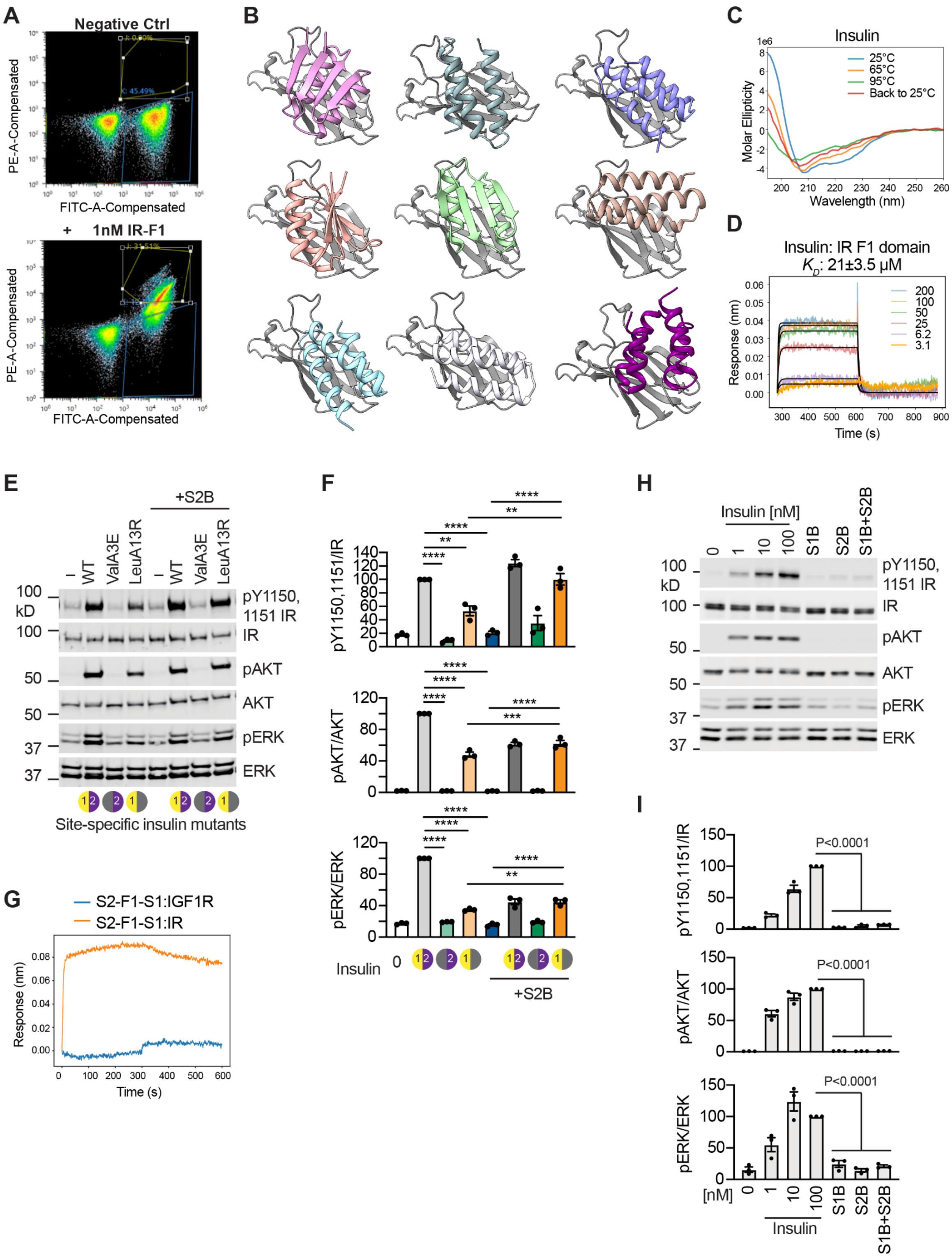

**Figure S1. Effects of site-1 binder and site-2 binder. Related to Fig2.**
(A) Yeast display of site-2 binder designs in presence of 1 nM of site-2 target protein (IR F1 domain).
(B) Representative structures of identified designed IR site-2 binders are shown. The IR F1 domain is depicted in gray ribbons, and the designed site-2 binders are shown in colored ribbons.
(C) CD spectra of insulin at various temperatures. Insulin unfolds as temperature increases and cannot be recovered when the temperature returns to room temperature (25 °C).
(D) Biolayer interferometry characterization of binding of Insulin to IR F1 domain. Global kinetic fit was reported.
(E) IR signaling in DKO-IR-A cells treated with 10 nM wild-type (WT) or site-specific insulin mutants with or without 100 nM site-2 binder (S2B) for 10 minutes.

(F) Quantification of the western blot data shown in (E). Levels of phosphorylation were normalized to total protein levels and shown as intensities relative to that in WT insulin alone. Mean ± sem. N=3 independent experiments. Significance calculated using two-tailed student's t-test.
(G) Biolayer interferometry characterization of binding of S2-F1-S1 to IR and IGF1R.
(H) IR signaling in DKO-IR-B cells treated with the indicated ligands for 10 minutes: 0, 1, 10, 100 nM insulin, 100 nM S1B, S2B or S1B+S2B.
(I) Quantification of the western blot data shown in (H). Levels of phosphorylation were normalized to total protein levels and shown as intensities relative to that in 100 nM insulin alone. Mean ± sem. N=3 independent experiments. Significance calculated using two-tailed student's t-test.

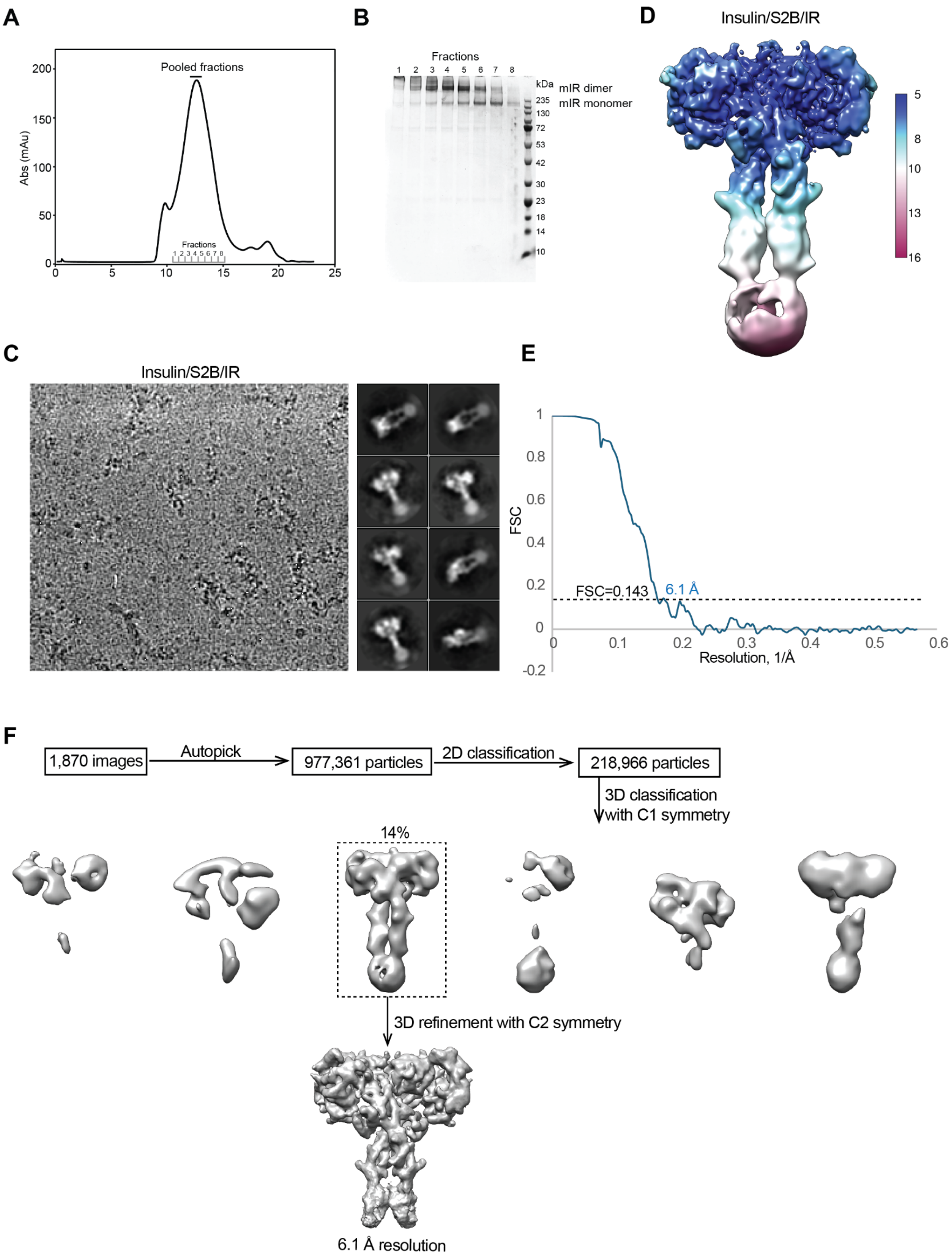

**Figure S2. Cryo-EM analysis of the Insulin/S2B/IR complex. Related to Figure. 2.**

(A) Representative size-exclusion chromatogram of mouse IR (mIR).

(B) The peak fractions in (A) were visualized on SDS-PAGE by Coomassie blue staining.

(C) Representative electron micrograph and 2D class averages of the insulin/S2B/IR complex.

(D) Unsharpened cryo-EM map colored by local resolution.

(E) The gold-standard Fourier Shell Correlation (FSC) curve for the cryo-EM map shown in Figure 2.

(F) Flowchart of cryo-EM data processing.

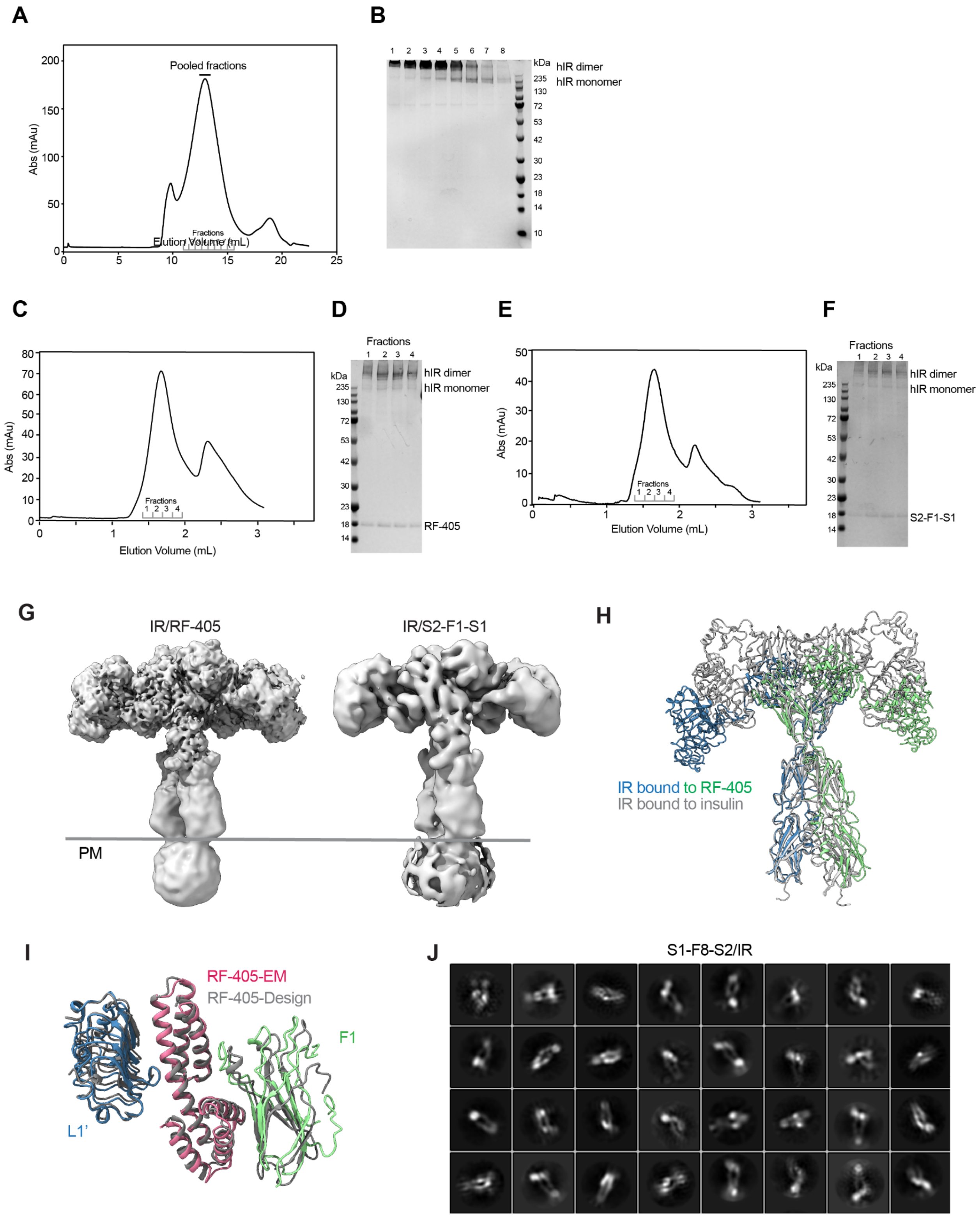

**Figure S3. Structures of RF-405/IR, S2-F1-S1/IR, and S1-F8-S2/IR. Related to Figure 4.**
(A) Representative size-exclusion chromatogram of human IR (hIR).
(B) The peak fractions in (A) were visualized on SDS-PAGE by Coomassie blue staining.
(C) Representative size-exclusion chromatogram of RF-405/hIR complex.
(D) The peak fractions in (C) were visualized on SDS-PAGE by Coomassie blue staining.
(E) Representative size-exclusion chromatogram of S2-F1-S1/hIR complex.
(F) The peak fractions in (E) were visualized on SDS-PAGE by Coomassie blue staining.
(G) Cryo-EM density of RF-405/IR and S2-F1-S1/IR.
(H) Overlay of the insulin/IR structure (PDB:6pxv, gray) with the RF-405/:IR complex structure.

(I) Overlay of the RF-405/IR design model (gray) with the RF-405/IR complex structure. The aligned binder RMSD is 1.06 Å.
(J) Representative 2D class averages of particles of S1-F8-S2/IR complex. S1-F8-S2 was not able to induce stable conformation and the particles were highly heterogeneous.

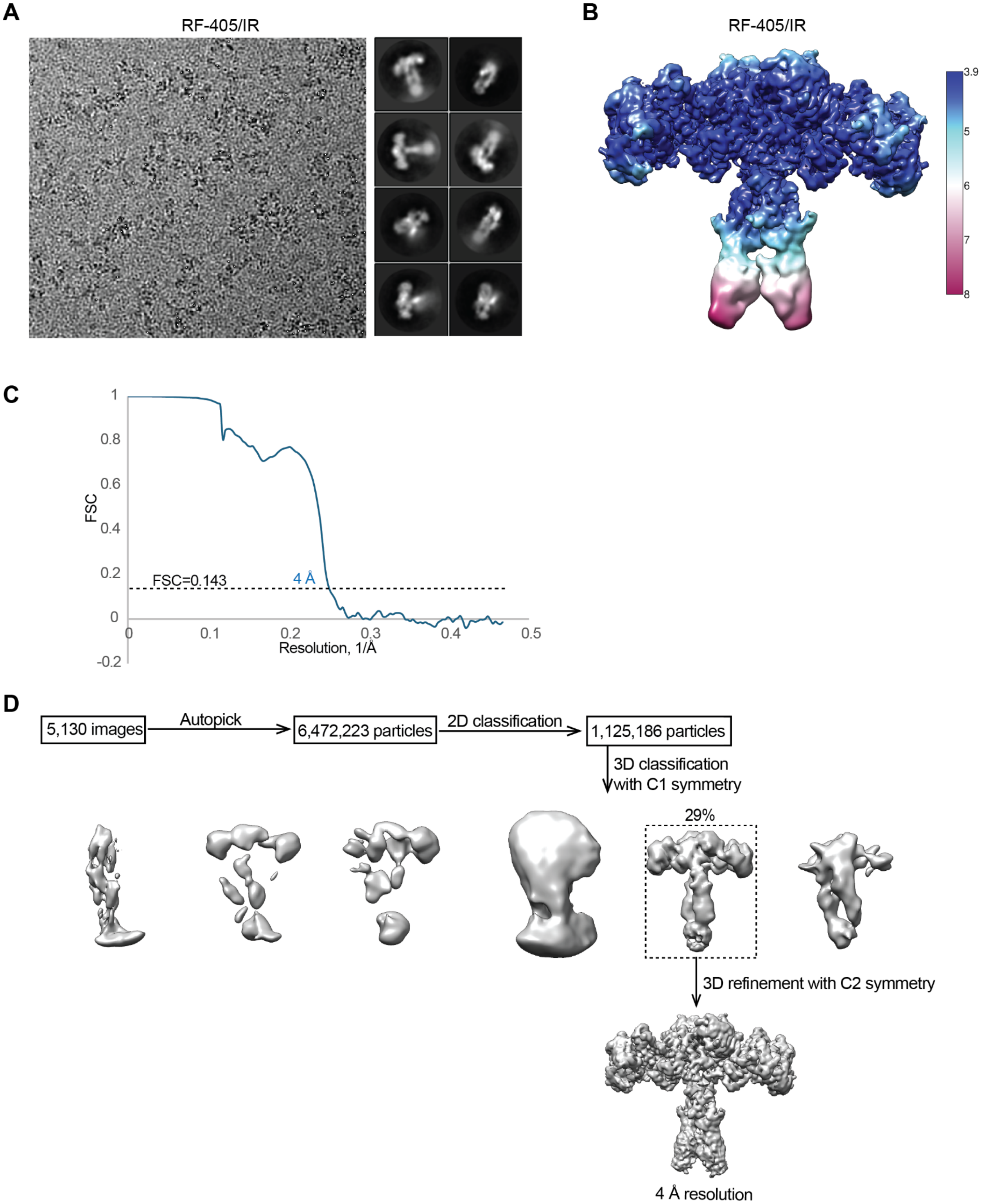

**Figure S4. Cryo-EM analysis of the RF-405/IR complex. Related to Figure. 4.**
(A) Representative electron micrograph and 2D class averages of the RF-405/IR complex.
(B) Unsharpened cryo-EM map colored by local resolution.
(C) The gold-standard Fourier Shell Correlation (FSC) curve for the cryo-EM map shown in Figure 4.
(D) Flowchart of cryo-EM data processing.

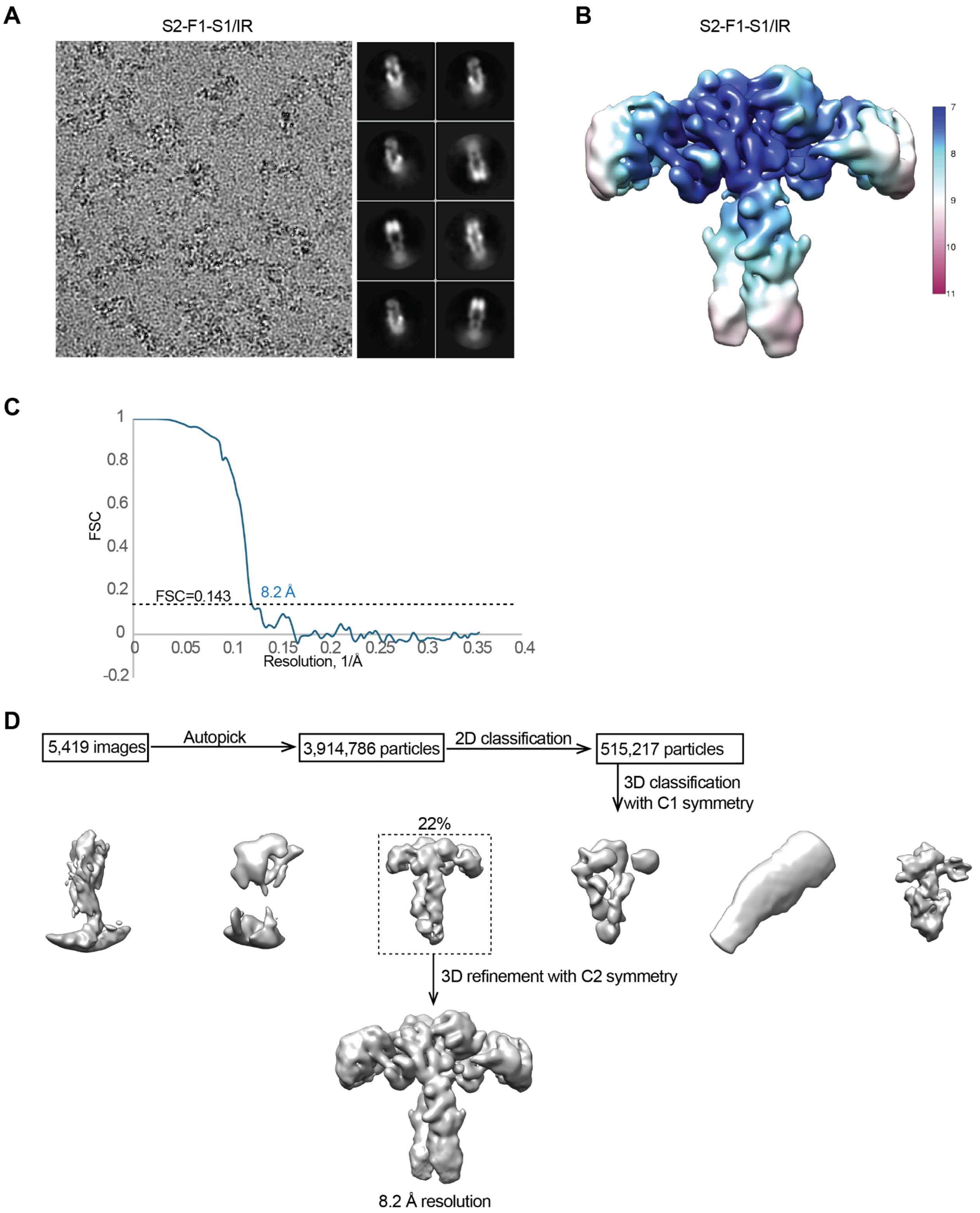

**Figure S5. Cryo-EM analysis of the S2-F1-S1/IR complex. Related to Figure. 4.**
(A) Representative electron micrograph and 2D class averages of the S2-F1-S1/IR complex.
(B) Unsharpened cryo-EM map colored by local resolution.
(C) The gold-standard Fourier Shell Correlation (FSC) curve for the cryo-EM map shown in Figure 4.
(D) Flowchart of cryo-EM data processing.

**A**

| Name | Linker Seq |
|---|---|
| S2-F5-S1 | GGSGG |
| S2-F3-S1 | GSG |
| S2-F2-S1 | GG |
| S2-F1-S1 | G |
| S1-F11-S2 | GGSGGSGGSGG |
| S1-F8-S2 | GGSGGSGG |
| S1-F5-S2 | GGSGG |
| S1-F3-S2 | GSG |

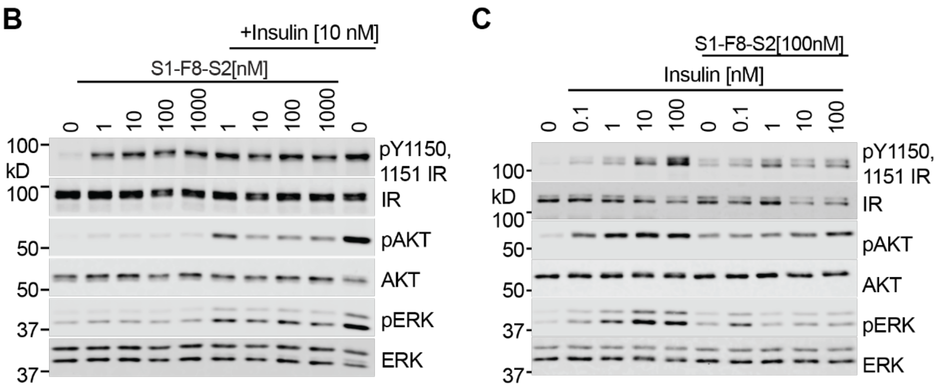

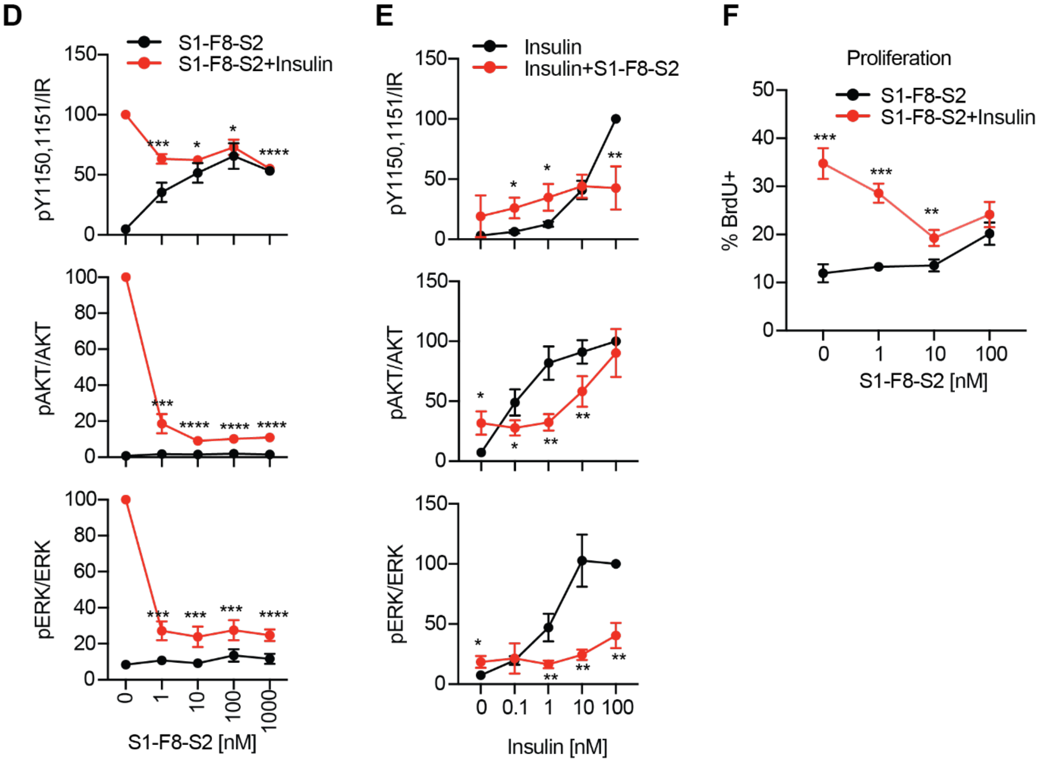

**G**

| IR intracellular domains | Insulin | RF-405 | RF-409 | S2-F1-S1 | S2-F5-S1 |
|---|---|---|---|---|---|
| | | | **Agonist, EC50 [nM]** | | |
| pY960 | 45.14 ± 1.35 | 11.94 ± 1.22**** | N.D | 77.76± 1.43** | N.D |
| pY1146 | 8.91 ± 1.37 | 7.13 ± 1.21 | N.D | 20.80 ± 1.36* | N.D |
| pY1150, pY1151 | 16.07 ± 1.38 | 7.29 ± 1.11**** | 8.23 ± 1.08** | 7.07 ± 1.19*** | 7.65 ± 1.55** |
| pY1316 IR | 23.70 ± 1.27 | 13.90 ± 1.20** | N.D | 86.96 ± 1.35**** | N.D |
| pY1322 IR | 41.60 ± 1.25 | 22.86 ± 1.22**** | N.D | 145.75 ± 1.44**** | N.D |
| pAKT | 2.16 ± 1.49 | 0.78 ± 1.38 | 1.24 ± 1.44 | 2.88 ± 1.40 | 9.20 ± 1.33 |
| pERK | 3.01 ± 1.15 | 4.72 ± 1.20 | 4.53 ± 1.20 | 96.89 ± 1.49*,[a] | 133.20 ± 1.45** |

**Figure S6. Antagonistic effects of S1-F8-S2. Related to Figure 5.**
(A) The sequence of linkers in flexibly-linked ligands.
(B) IR signaling in DKO-IR-B cells by insulin and S1-F8-S2. Cells were treated with the indicated concentrations of S1-F8-S2 for 1 hour and then treated with 10 nM insulin for 10 minutes.
(C) IR signaling in C2C12-IR cells by insulin and S1-F8-S2. Cells were treated with 100 nM S1-F8-S2 for 1 hour and then treated with the indicated concentrations of insulin for 10 minutes.
(D) Quantification of the western blot data shown in (B). Levels of phosphorylation were normalized to total IR levels and shown as intensities relative to that in WT insulin alone. Mean ± SD. N=3 independent experiments. Significance calculated using 2-way ANOVA. *p<0.05, ***p<0.001, and ****p<0.0001.

(E) Quantification of the western blot data shown in (C). Levels of phosphorylation were normalized to total IR levels and shown as intensities relative to that in WT insulin alone. Mean ± SD. N=4 independent experiments. Significance calculated using 2-way ANOVA. *$p<0.05$ and $p<0.01$.
(F) Cell proliferation. C2C12-IR cells were incubated with the indicated concentrations of S1-F8-S2 for 24 hours in the presence or absence of 10 nM insulin. Cells were then incorporated with Bromodeoxyuridine (BrdU) for 2 hours. BrdU-positive cells were analyzed by FACS. Mean ± SD. N= 4 independent experiments. Significance calculated using 2-way ANOVA. **$p<0.01$ and ***$p<0.001$.
(G) Summary of EC50. Mean ± sem. EC50 values obtained from dose-response curves in Fig. 5C-I. p values were calculated by Extra sum-of-squares F Test in Prism, between insulin and ligands. p values vs insulin. *$p<0.05$, **$p<0.01$, ***$p<0.001$, and ****$p<0.0001$. p value vs S2-F5-S1, [a]$p<0.05$. ND, not determined.

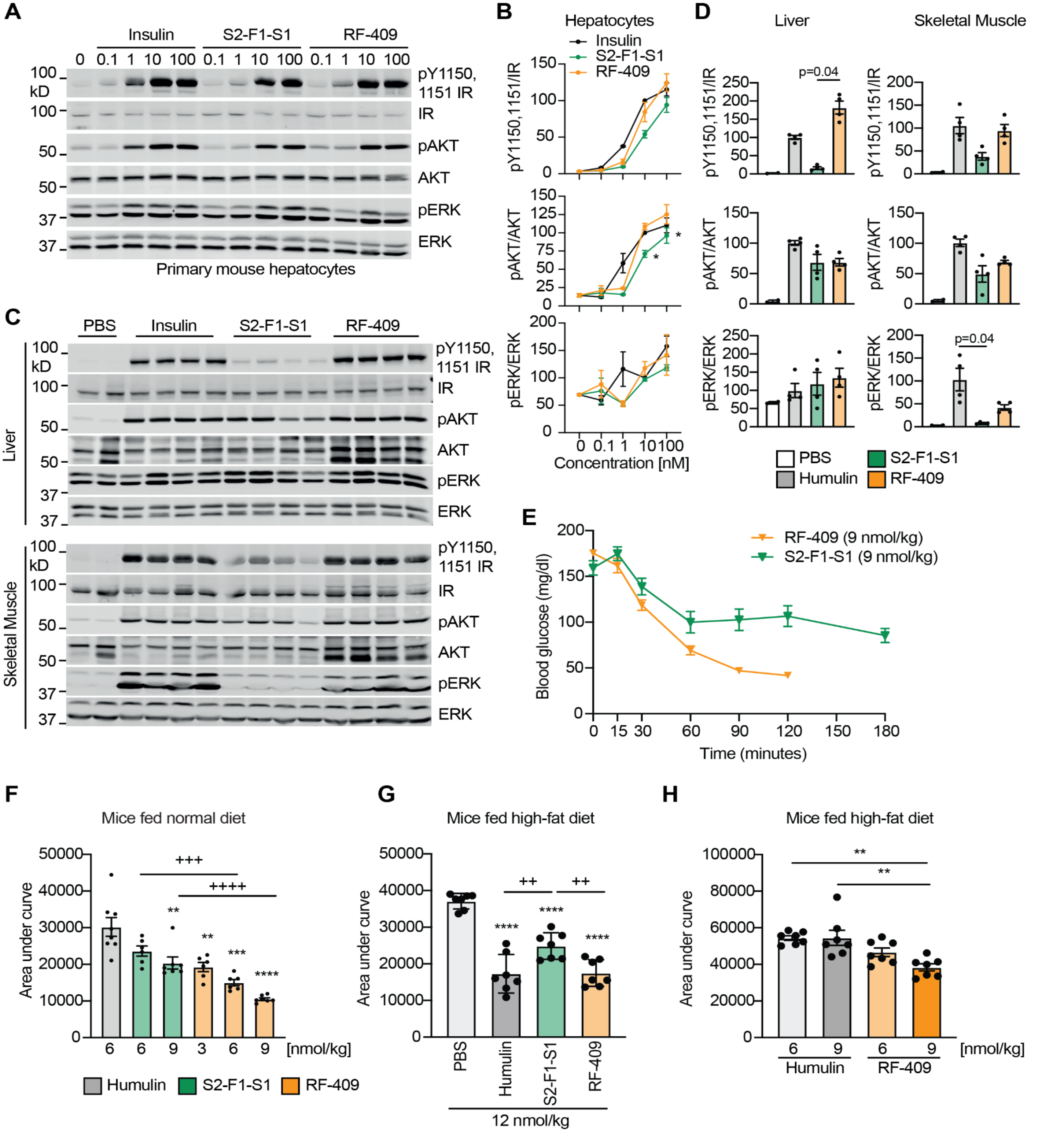

**Figure S7. Designed IR agonists activate IR signaling and control glucose levels in mice. Related to Figure 7.**

(A) IR signaling in primary mouse hepatocytes Treated with the indicated ligands for 10 minutes.

(B) Quantification of the western blot data shown in (A). Levels of phosphorylation were normalized to total protein levels and shown as intensities relative to that in WT insulin. Mean ± SD. N = 3 independent experiments. Significance calculated using 2-way ANOVA. p values vs insulin. *p<0.05.

(C) IR signaling in the liver and skeletal muscle of mice. Each lane contains lysate from an individual mouse.

(D) Quantification of data in (C). Levels of protein phosphorylation were normalized to total protein levels and shown as intensities relative to that in insulin-treated conditions. Mean ± sem. N= 4 mice per group. PBS, N=2. Significance calculated using 1-way ANOVA.

(E) Insulin tolerance test in mice fed normal chow diet. Mice were injected intraperitoneally with S2-F1-S1 or RF-409 at the indicated doses, and their blood glucose levels measured at the indicated time points after injection. Mean ± sem. N= 7 mice per group.
(F) Glucose area under the curve during ITT in **Fig. 7B** and **Fig. S7E**. Significance calculated using two-tailed student's t-test.
(G) Glucose area under the curve during ITT in **Fig. 7C**. Significance calculated using two-tailed student's t-test.
(H) Glucose area under the curve during ITT in **Fig. 7D**. Significance calculated using two-tailed student's t-test.

**Table S1. Cryo-EM data collection and structure refinement statistics.**

| Structure | Insulin/S2B/IR<br>EMD-47043<br>PDB: 9DNN | RF-405/IR<br>EMD-47031<br>PDB: 9DN6 | S2-F1-S1/IR<br>EMD-47041<br>PDB: 9DNI |
|---|---|---|---|
| Magnification | 45,000 | 130,000 | 81,000 |
| Voltage (kV) | 200 | 300 | 300 |
| Electron exposure ($e^-/Å^2$) | 60 | 60 | 60 |
| Defocus range (μm) | 1.2-2.2 | 1.2-2.2 | 1.2-2.2 |
| Pixel size (Å) | 0.88 | 1.07 | 1.404 |
| Symmetry imposed | C2 | C2 | C2 |
| Initial particle images (no.) | 977,361 | 6,472,223 | 3,914,786 |
| Final particle images (no.) | 10,481 | 66,409 | 23,881 |
| Map resolution (Å) | 6.1 | 4.0 | 8.2 |
| FSC threshold | 0.143 | 0.143 | 0.143 |
| Initial model used (PDB code) | 6PXV | 6PXV | 6PXV |
| Model Composition | | | |
| Non-hydrogen atoms | 29,774 | 28,535 | 28,144 |
| Protein residues | 1,866 | 1,766 | 1,742 |
| R.m.s. deviations | | | |
| Bond length (Å) | 0.003 | 0.004 | 0.004 |
| Bond angle (°) | 0.718 | 0.678 | 0.676 |
| Validation | | | |
| Molprobity score | 2.5 | 2.24 | 2.49 |
| Clashscore | 27.22 | 13.18 | 26.9 |
| Poor rotamers (%) | 0 | 0.44 | 0 |
| Ramachandran plot | | | |
| Favored (%) | 89.05 | 87.46 | 89.01 |
| Allowed (%) | 10.9 | 12.14 | 10.87 |
| Outliers (%) | 0.05 | 0.4 | 0.12 |
